# Altered calcium signaling in Bergmann glia contributes to Spinocerebellar ataxia type-1

**DOI:** 10.1101/2023.05.09.539932

**Authors:** Jose Antonio Noriega-Prieto, Carmen Nanclares, Francisco E. Labrada-Moncada, Marija Cvetanovic, Alfonso Araque, Paulo Kofuji

## Abstract

Spinocerebellar ataxia type 1 (SCA1) is a neurodegenerative disease caused by an abnormal expansion of glutamine (Q) encoding CAG repeats in the *ATAXIN1* (*ATXN1*) gene and characterized by progressive cerebellar ataxia, dysarthria, and eventual deterioration of bulbar functions. SCA1 shows severe degeneration of cerebellar Purkinje cells (PCs) and activation of Bergmann glia (BG), a type of cerebellar astroglia closely associated with PCs. Using electrophysiological recordings, confocal imaging and chemogenetic approaches, we have investigated the electrical intrinsic and synaptic properties of PCs and the physiological properties of BG in SCA1 mouse model expressing mutant ATXN1 only in PCs. PCs of SCA1 mice displayed lower spontaneous firing rate and larger medium and slow afterhyperpolarization currents (mI_AHP_ and sI_AHP_) than wildtype mice, whereas the properties of the synaptic inputs were unaffected. BG of SCA1 mice showed higher calcium hyperactivity and gliotransmission, manifested higher frequency of NMDAR-mediated slow inward currents (SICs) in PC. Preventing the BG calcium hyperexcitability of SCA1 mice by loading BG with the calcium chelator BAPTA restored mI_AHP_ and sI_AHP_ and spontaneous firing rate of PCs to similar levels of wildtype mice. Moreover, mimicking the BG hyperactivity by activating BG expressing Gq-DREADDs in wildtype mice reproduced the SCA1 pathological phenotype of PCs, i.e., enhancement of mI_AHP_ and sI_AHP_ and decrease of spontaneous firing rate. These results indicate that the intrinsic electrical properties of PCs, but not their synaptic properties, were altered in SCA1 mice, and that these alterations were associated with the hyperexcitability of BG. Moreover, preventing BG hyperexcitability in SCA1 mice and promoting BG hyperexcitability in wildtype mice prevented and mimicked, respectively, the pathological electrophysiological phenotype of PCs. Therefore, BG plays a relevant role in the dysfunction of the electrical intrinsic properties of PCs in SCA1 mice, suggesting that they may serve as potential targets for therapeutic approaches to treat the spinocerebellar ataxia type 1.

## Introduction

Spinocerebellar ataxia type 1 (SCA1) belongs to the group of the spinocerebellar ataxias (SCAs), a large and diverse group of autosomal-dominant, progressive neurodegenerative diseases. The most common SCAs are caused by the expansion of a CAG nucleotide repeat that encodes polyglutamine (polyQ) in the pertinent disease proteins. In the case of SCA1 the expanded polyQ sequences are found in ataxin 1 (ATXN1) protein (Paulson et al., 2017). While the physiological function of ataxin-1 is not completely understood, it appears to be involved in regulating gene expression (Serra et al. 2004, Ingram et al,2016, Cvetanovic 2011, Handler 2023). ATXN1 containing a wild-type polyQ tract shuttles between the cytoplasm and nucleus. SCA1 is characterized by progressive motor deficits, cognitive decline, and mood changes, with longer number of CAG repeats correlating with disease severity.

A hallmark of SCA1 is the loss of cerebellar Purkinje neurons (Nino et al., 1980; Burright et al., 1995; Servadio et al., 1995; Seidel et al., 2012). This well-defined attribute has led to numerous studies to focus on neuronal cells to understand the molecular mechanisms underlying the disease. However, accumulating evidence indicate that glial cells are prominent contributors to many neurodegenerative diseases, such as Alzheimeŕs disease (Phillips et al., 2014; Fakhoury, 2018; Nanclares et al., 2021), Huntingtońs disease (Shin et al., 2005; Jiang et al., 2016; Meunier et al., 2016), Parkinsońs disease (Halliday and Stevens, 2011; Díaz et al., 2019), amyotrophic lateral sclerosis (Yamanaka et al., 2008), and some forms of ataxia (Kretzschmar et al., 2005; Custer et al., 2006; Furrer et al., 2011), including SCA1 (Kim et al 2018). Astroglial cells play key roles in numerous functions within the brain, such as regulation of regional blood flow, ionic balance of the extracellular space, and modulation of synaptic transmission (Kofuji and Newman, 2004; Perea et al., 2014; Sloan and Barres, 2014). The cerebellar cortex contains three main types of astroglial cells: Bergmann glia (BG), a specialized type of cerebellar astrocytes, protoplasmic velate astrocytes, and white matter fibrous astrocytes (Araujo et al., 2019). BG somas, located in the Purkinje cell layer, extend their processes along the dendrites of Purkinje neurons into the molecular layer, where they envelop and regulate excitatory synapses on Purkinje neurons (Yamada and Watanabe, 2002; Sasaki et al., 2012; Buffo and Rossi, 2013; De Zeeuw and Hoogland, 2015). Because of their location and close interactions, BG are crucial for the function of Purkinje neurons. Alterations in BG function through altered calcium signaling or potassium and glutamate uptake, for example, can lead to dysfunction of adjacent Purkinje neurons (Kim et al., 2018).

Present study has investigated the potential alteration of the functional properties of BG and PCs in a SCA1 mouse model, and their interaction focused our interest on possible changes that may be occurring on PC and BG synaptic inputs as well as their intrinsic properties in SCA1. Accordingly, we aimed to decipher the contribution that BG dysregulation has on PC pathogenesis in SCA1. Our results show that the cellular phenotype of SCA1 mice was manifested as a reduced spontaneous firing rate of PCs and an enhancement of mIAHP and sIAHP, which were accompanied by a calcium hyperactivity of BG. Moreover, preventing the BG calcium activity in SCA1 mice, restored the normal phenotypic properties of the spontaneous firing rate of PCs and mIAHP and sIAHP, as observed in WT mice. Finally, stimulating the BG calcium activity in WT recapitulated the cellular phenotype of SCA1 mice. Therefore, BG decisively contributes to the neuronal phenotype of the SCA1 neuropathology.

## Materials and Methods

### Animals

The creation of the *ATXN1[82Q]* mice was previously described (Burright et al., 1995). The murine Pcp2 (L7) promoter was used to direct the expression of the human SCA1-coding region containing an expanded allele containing 82 repeats. Thus, this mouse model will express mutant ATXN1 only in PCs. *ATXN1[82Q]* mice originally on FVB background were gifts from Dr. Harry Orr and were backcrossed onto a C57/Bl6 background. *ATXN1[82Q]* and control littermates were used from 2- to 8-month-old. Efforts were made to make the groups equal between males and females. Mice were housed in a temperature-and humidity-controlled room on a 12 h light/12 h dark cycle with access to food and water *ad libitum*. All animal care and sacrifice procedures were approved by the University of Minnesota Institutional Animal Care and Use Committee (IACUC) in compliance with the National Institutes of Health guidelines for the care and use of laboratory animals.

### Slice electrophysiology

Acute sagittal cerebellar slices (250 μm thick) were prepared in an ice-cold high sucrose solution of the following composition (in mM): sucrose 189, glucose 10, NaHCO_3_ 26, KCl 3, MgSO_4_ 5, CaCl_2_ 0.1, NaH_2_PO_4_ 1.25. Afterward, slices were incubated at room temperature in artificial cerebrospinal fluid (ACSF) for 45 min-1 hr before use. ACSF used for ex vivo experiments contained in mM: NaCl 124, KCl 5, NaH_2_PO_4_ 1.25, MgSO_4_ 2, NaHCO_3_ 26, CaCl_2_ 2, glucose 10; gassed with 95% O_2_, 5% CO_2_. The pH was adjusted to 7.4 with NaOH. For studies, slices were transferred to an immersion recording chamber, superfused at 2 mL/min with oxygenated ACSF, and visualized under an Olympus BX50WI microscope (Olympus Optical; Japan).

#### Electrophysiology in Purkinje cells

We visually identified Purkinje cells based on size and location between the granule cell layer and the molecular layer. The following whole-cell recordings in PCs were performed. Slow inward current recordings (SICs) were performed in ACSF of the following composition (in mM): NaCl 124, KCl 2.69, KH_2_PO_4_ 1.25, NaHCO_3_ 26, CaCl_2_ 4, glucose 10, and glycine 10 μM. Magnesium was not included in the solution to optimize the activation of NMDA receptors. Additionally, 1 μM of tetrodotoxin (TTX) was included. SICs were defined as currents with a rise time > 5 ms, decay time > 10 ms and ΤonL>L5. Input-Output curves of evoked excitatory postsynaptic currents (EPSCs) and paired-pulse ratio were recorded in ACSF with picrotoxin 50 μM and CGP54626 1 µM to block GABA-A and GABA-B inhibitory neurotransmission, respectively. To measure climbing fiber synaptic inputs CNQX 500 nM was added to reduce the size of synaptic currents (Tsai et al., 2012). The internal pipette solution for these experiments contained (in mM): Cs-gluconate 117, HEPES 20, EGTA 0.4, NaCl 2.8, TEA-Cl 5, Mg-ATP 2, Na-GTP 0.3. The pH was adjusted to 7.3 with CsOH. Spontaneous and evoked action potential firing was recorded in ACSF alone. Spontaneous action potential firing was recorded using the voltage-clamp mode of the cell-attached configuration. The internal pipette solution for spontaneous firing rate contained the same composition as the external solution and in the case of evoked firing rate the internal solution contained (in mM): K-gluconate 135, KCl 10, HEPES 10, MgCl 1, Mg-ATP 2. The pH was adjusted to 7.3 with KOH. The medium and slow afterhyperpolarization currents (mI_AHP_ and sI_AHP_ respectively) were studied using a hyperpolarizing pulse of 800 ms duration. Membrane potential was held at −70 mV and a depolarized pulse at 0 mV during 800 ms was applied. While the mI_AHP_ was measured as the peak amplitude of the afterhyperpolarization current, the sI_AHP_ was estimated as the area under the curve during 1 s period beginning at the peak of the current (Santini et al., 2008; Maglio et al., 2021). The internal pipette solution for I_AHP_ was the same one used for evoked firing rate.

Recordings were obtained from lobules V and VI Purkinje cells (PCs) with PC-ONE amplifiers (Dagan Instruments; Minneapolis, MN). Membrane potentials were held at −70 mV unless otherwise stated. Signals were filtered at 1 kHz, acquired at a 10 kHz sampling rate, and fed to a Digidata 1440A digitizer (Molecular Devices; San Jose, CA). pCLAMP 10.4 (Axon Instruments, Molecular Devices; San Jose, CA) was used for stimulus generation, data display, data acquisition, and data storage. To record evoked excitatory postsynaptic currents (EPSCs), theta capillaries filled with ACSF were used for bipolar stimulation and placed in the vicinity of the PC. Input-output curves were made by increasing stimulus intensities from 0 to 100 μA. Paired pulse ratio was done by applying paired pulses (2 msec duration) with 25, 50, 75, 100, 150, 200, 300, and 500 ms inter-pulse intervals. The paired-pulse ratio was calculated by dividing the amplitude of the second EPSC by the first (PPR = EPSC-2 / EPSC-1). Action potentials were evoked by the injection of current (from −50 to 1400 pA in 50 pA steps) through the electrode inside the patch pipette. The amplitude of the synaptic inputs as well as the frequency of evoked action potential (Hz = spike/s) were analyzed using Clampfit 10.4 (Axon Instruments, Molecular Devices). Spontaneous action potentials were recorded for 5 min in the cell-attached mode. Spike sorting and analysis were performed with Spike2 (Cambridge Electronic Design Limited, CED). Spike firing frequency (Hz = spike/s) and firing pattern variability were calculated for each Purkinje cell. Variability in firing pattern was quantified using the coefficient of variation (CV) of the interspike time interval (ISI) [CV = (standard deviation of ISIs) / (mean of ISIs)]. To measure rhythmicity during burst firing, CV2 of the ISI was calculated [CV2 = 2(| ISI_After_ – ISI_Before_ |) / (ISI_After_ + ISI_Before_), where for each given spike, ISI_After_ is the ISI to the next spike and ISI_Before_ is the ISI to the previous spike. CV2 is calculated for every spike and then the mean CV2 is determined (Holt et al., 1996). All electrophysiological recordings were performed at 30-32 °C.

#### Electrophysiology in Bergmann glia

BG were identified based on a combination of morphological and electrical properties. They are localized in the PC layer and have small cell round bodies. Biophysically, astrocytes are characterized by a low input resistance, they do not fire action potentials and typically rest at very hyperpolarized potentials (approximately −70/-80mV, close to the potassium equilibrium potential). Thus, once entered in the whole-cell mode an application of several square electrical pulses (from −120 to 40 mV at 10 mV steps) was delivered. Under these conditions, astrocytes display a quasi-linear voltage-current relationship and as stated above they do not fire action potentials. To evoke glutamate transporter currents theta capillaries filled with ACSF were used for bipolar stimulation and placed in the surroundings of the BG patched. Neuronal terminals were stimulated during 500 ms at 75 Hz. To study the K^+^ buffer capacity of BG we measured Kir4.1 currents. A pre-pulse to 0 mV during 300 ms was applied followed by 500 ms of increasing voltages (from −160 to +60 mV in 10 mV steps). The first depolarizing step serves to remove all transient outward potassium current components, which will undergo steady-state inactivation at this potential. Once a stable current is obtained BaCl_2_ 100 µM is perfused and the same protocol is applied until it is stabilized again. Ba^2+^ is a nonspecific potassium channel inhibitor, but the low concentration used in these experiments is thought to be specific for Kir channels. Then, subtracting the I-V traces we isolated Ba^2+^–sensitive Kir4.1 currents (Milner, 2012).

In some experiments, BG was recorded with a patch-clamp pipette filled with 20 mM BAPTA (Ca^2+^ chelator). BAPTA was allowed to diffuse for at least 30 min. It has been previously described that BG are electrically coupled through gap junctions (Müller et al., 1996). Muller et al loaded a BG through the patch pipette with the dye Lucifer Yellow and described that after 30-40 min the dye diffused among about 25 BG coupled cells. Considering that the size of a BG soma ranges around 7-10 µm, that would make a total distance of dye diffusion of 175-250 µm. To make sure that the BAPTA had diffused in our recordings we restricted our study to paired-recorded PC and BG in relatively close proximity (<120 µm).

### In vitro Ca^2+^ imaging

Ca^2+^ levels in Bergmann glia from lobules V/VI were monitored using a Leica SP5 multiphoton microscope (model DM6000 CFS upright multiphoton microscope with TCS SP5 MP laser, Leica). Ca^2+^ was monitored using tamoxifen-induced GCaMP6 transgenic mice, Cre-mediated targeted BG as Ca^2+^ sensor. Cells were illuminated every 100 ms at 490 nm with a camera control and synchronization for the two-photon imaging by LAS software (Leica). All Ca^2+^ experiments were performed in the presence of TTX (1 μM).

Videos were obtained at 512□×□512 resolution with a sampling interval of 1 s. A custom MATLAB program (Calsee: https://www.araquelab.com/code/) was used to quantify fluorescence levels in BG. Fluorescent Ca^2+^ variation frequency recorded from the cells was given as the number of events per minute (min^-1^) where BG Ca^2+^ signal was quantified from the spontaneous Ca^2+^ event frequency, which was calculated from the number of Ca^2+^ elevations over 5 minutes of video recording.

To determine the BG responding to ATP puff, Ca^2+^ event probability was calculated, in which the number of Ca^2+^ elevations was grouped in 5-s bins, and values of 0 and 1 were assigned for bins showing either no response or a Ca^2+^ event, respectively. The Ca^2+^ event probability was calculated by dividing the number of regions of interest (ROIs) showing an event at each time bin by the total number of monitored ROIs (Navarrete and Araque, 2010). The Ca^2+^ event frequency was calculated for each slice, and for statistical analysis, the sample size corresponded to the number of slices as different slices were considered as independent variables.

### Viral injections

Mice were given injections of either DREADDSs (AAV8-GFAP-hM3D(Gq)-mCherry, UMN vector core) or control virus (AAV8-GFAP-mCherry, UMN vector core) into the lobule VI region of the cerebellum (AP: −6.6, ML: ±1, and DV: −1.6 from Bregma) and were delivered at a rate of 1 nL/sec for a total of 1000 nL using a Hamilton syringe and micropump controller (World Precisions Instruments). The syringes were left in place for 10 minutes after injection before being slowly withdrawn. The injections were administered bilaterally. DREADDs activation experiments were not conducted until at least 3 weeks after the surgery to ensure sufficient time for the viral expression.

## Results

### Electrophysiological properties of Purkinje cells in SCA1 mice

#### Spontaneous firing rate of PC is impaired in SCA1 mice

To characterize the functional properties of PC in SCA1 mice, we first assessed the spontaneous action potential firing rates of PCs of WT and SCA1 mice. Electrophysiological recordings of PCs in lobules V/VI (Figure 1A) were performed using the cell-attached configuration of the patch-clamp technique.

**Figure 1.**
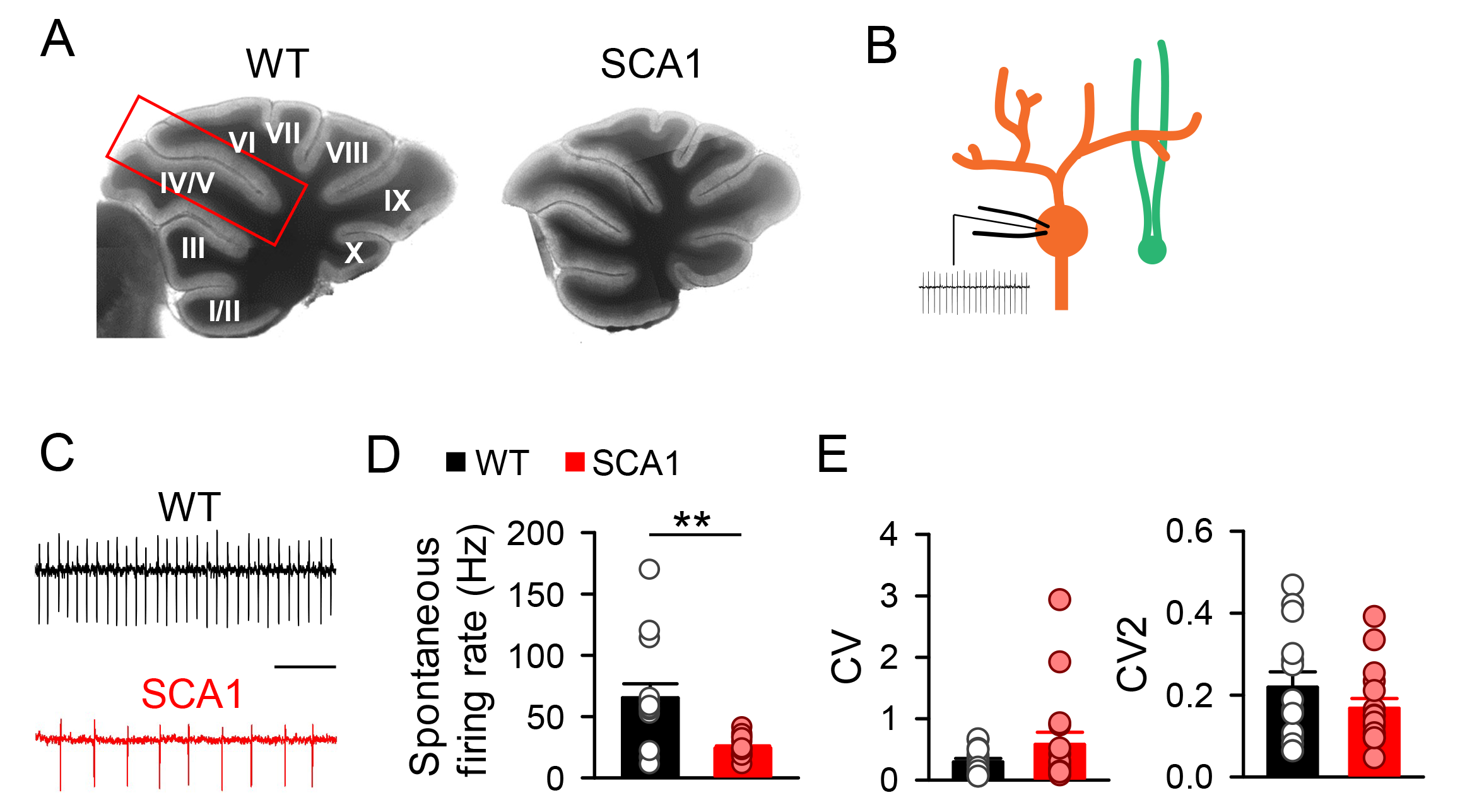
Purkinje cell spontaneous firing rate is decreased in SCA1 mouse model. **A.** Differential interference contrast (DIC) image of a cerebellum brain slice from WT (left) and SCA1 (right) mice. **B.** Schematic representation of spontaneous firing recording (black trace) from a PC (orange cell) close to a BG cell (green cell). **C.** Cell-attached recordings of spontaneous action potential firing in PCs from WT (black) and SCA1 (red) mice (scale bar: 100ms). **D.** Quantitative analysis of PC spontaneous firing rate (WT, *n* = 14 cells; SCA1, *n* = 16 cells; Mann-Whitney test; p=0.003). Note the lower firing rate in SCA1. **E.** Coefficient of variation (CV, left) and coefficient of variation 2 (CV2, right) of the same cells depicted in B (WT, *n* = 14 cells; SCA1, *n* = 16 cells; Mann-Whitney test CV p= 0.418, CV2 p= 0.418).

The spontaneous firing rate of PCs in SCA1 mice was significantly lower than in WT mice (Figure 1B, C and D). Moreover, because specific spatiotemporal patterns of spiking activities and silent intervals encodes information in the cerebellum (De Zeeuw et al., 2011), we further analyzed the coefficients of variation (CV and CV2) of the interspike intervals to quantify the firing regularity. Small values close to 0 indicate regular firing, whereas large values close to or >1 indicate irregular firing (Holt et al., 1996). There were no statistical differences in either CV or CV2 of the interspike intervals of PCs in WT and SCA1 mice (Figure 1E), indicating similar spontaneous firing regularity in both mice.

#### Excitatory synaptic transmission is not altered in SCA1 mice

The reduced spontaneous firing rate of PCs in SCA1 mice could be due to alterations of the excitatory synaptic input or to changes in the intrinsic electrical properties that determine the neuronal excitability. We first studied the synaptic properties of the two main excitatory synaptic inputs to PCs, i.e., synapses of climbing (CFs) and parallel (PFs) fibers (Konnerth et al., 1990) (Figure 2A).

**Figure 2.**
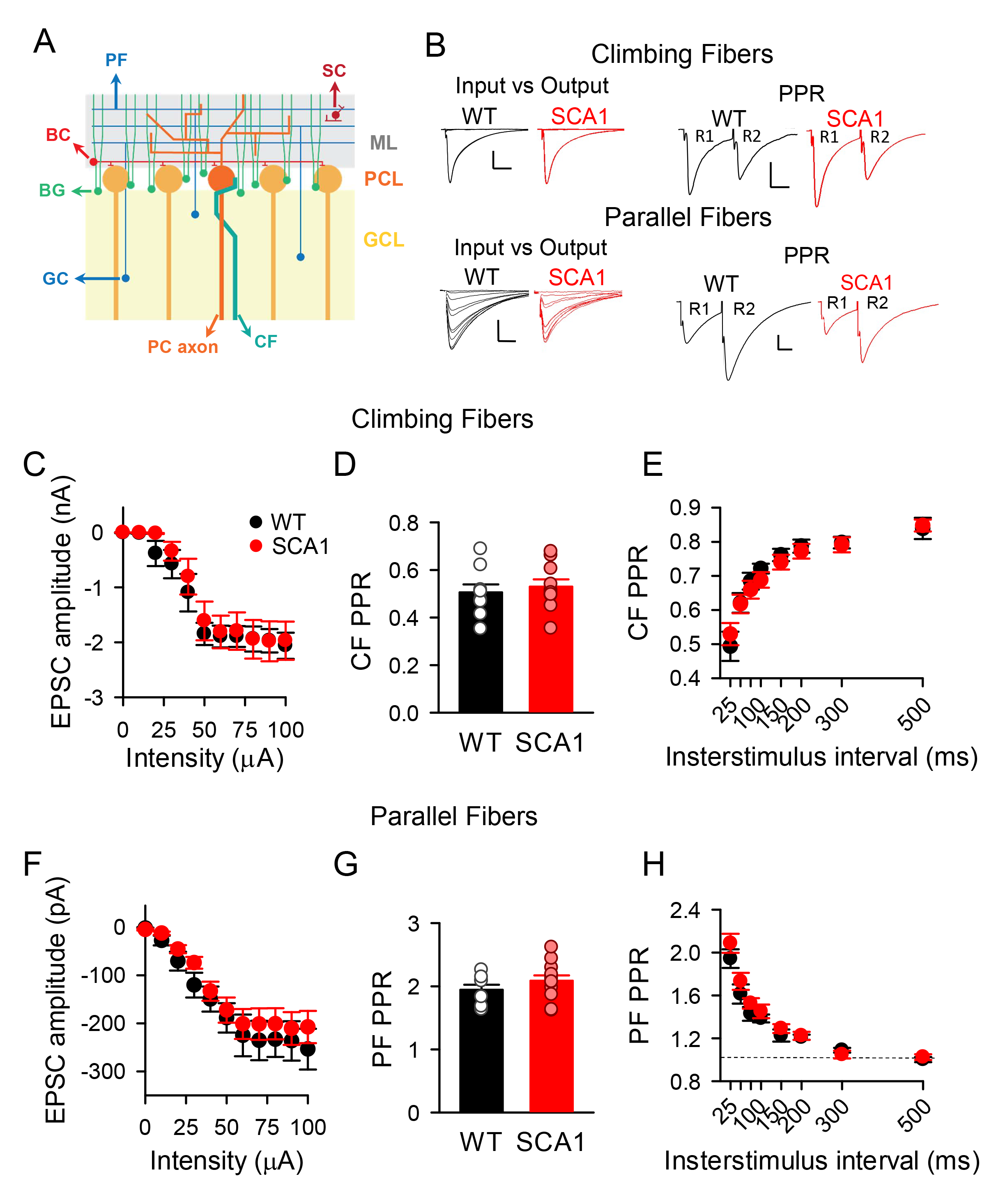
Excitatory synaptic transmission onto Purkinje cell is not altered in SCA1 mouse model. **A.** Schematic representation of synaptic inputs onto PCs (ML: Molecular layer, PCL: Purkinje cell layer, GCL: granule cell layer, GC: granule cells, PC: Purkinje cell, BC: Basket cells, SC: stellate cells, PF: parallel fiber, CF: climbing fiber). **B.** Representative traces of excitatory postsynaptic currents (EPSCs) evoked by climbing fiber stimulation (Top). Input-Output relationships of EPSCs (left, scale bar: 1 nA, 10 ms). Paired-pulse ratio (PPR) induced by two consecutive stimuli delivered at 25-ms interval (right, scale bar: 1 nA, 10 ms). Representative traces of excitatory postsynaptic currents (EPSCs) evoked by parallel fiber stimulation (Bottom). Input-Output relationships of EPSCs (left, scale bar: 100 pA, 10 ms). Paired-pulse ratio (PPR) induced by two consecutive stimuli delivered at 25-ms interval (right, scale bar: 100 pA, 10 ms). **C.** Input-Output relationships of excitatory postsynaptic currents (EPSCs) evoked by climbing fiber stimulation (WT, n = 8 cells; SCA1, n = 9 cells; two-way ANOVA with post hoc Tukey comparisons). **D.** Paired-pulse ratio (PPR) induced by two consecutive stimuli delivered at 25-ms interval. EPSCs were evoked by climbing fiber stimulation (WT, *n* = 9 cells; SCA1, *n* = 10 cells; t-test, two-tailed *p=*0.609). **E.** PPR induced by two consecutive stimuli delivered at different time intervals. EPSCs were evoked by climbing fiber stimulation (WT, n = 7 cells; SCA1, n = 10 cells; two-way ANOVA with post hoc Tukey comparisons). **F.** Input-Output relationships of EPSCs evoked by parallel fiber stimulation (WT, n = 8 cells; SCA1, n = 12 cells; two-way ANOVA with post hoc Tukey comparisons). **G.** PPR induced by two consecutive stimuli delivered at 25-ms interval. EPSCs were evoked by parallel fiber stimulation (WT, *n* = 8 cells; SCA1, *n* = 12 cells; t-test, two-tailed *p=*0.264). **H.** PPR induced by two consecutive stimuli delivered at different time intervals. EPSCs were evoked by parallel fiber stimulation (WT, n = 8 cells; SCA1, n = 12 cells; two-way ANOVA with post hoc Tukey comparisons).

We performed whole-cell electrophysiological recordings of PCs in lobules V/VI and stimulated either CFs or PFs using an extracellular electrode located in the granule cell or molecular layer. Consistent with previous reports (Konnerth et al., 1990; Liu et al., 2022), CF stimulation elicited excitatory postsynaptic currents (EPSCs) characterized by their large amplitude (> 500 pA), their all-or-none responses and depression of the paired-pulse ratio (PPR), whereas PF stimulation evoked EPSCs of modest amplitude (in the range of a few hundred pA) and facilitation of the PPR. The CF-or PF-evoked EPSC amplitudes (Figure 2B), their input-output curves (Figure 2C and F), and their PPR (Figure 2D, E, G and H) were not significantly different in WT and SCA1 mice, indicating that the synaptic properties of the excitatory synaptic inputs to PCs were not altered in SCA1 mice.

#### Intrinsic electrical properties of PCs are altered in SCA1 mice

Since the synaptic properties were not affected in SCA1 mice, we then investigated whether the alternative mechanism potentially involved in the reduced spontaneous firing rate of PCs, i.e., the intrinsic neuronal electrical properties, were altered in SCA1 mice. We first studied the intrinsic electrical excitability of PCs, quantifying the number of action potentials evoked by 1 s depolarizing current pulses injected into PCs. The evoked firing rate initially increased as the depolarizing pulse intensity increased, reached a maximum and then decreased as the depolarization further increased, through a phenomenon called spike-frequency adaptation, i.e., a reduction in the firing frequency of their spike response following an initial increase (Figure 3A, B). No significant differences between WT and SCA1 mice were found in the increasing phase and the maximum values of the firing rate, whereas a stronger adaptation in the decreasing phase occurred in SCA1 mice compared with WT mice (Figure 3A, B), indicating that the intrinsic electrical properties of PC were altered in the SCA1 mice.

**Figure 3.**
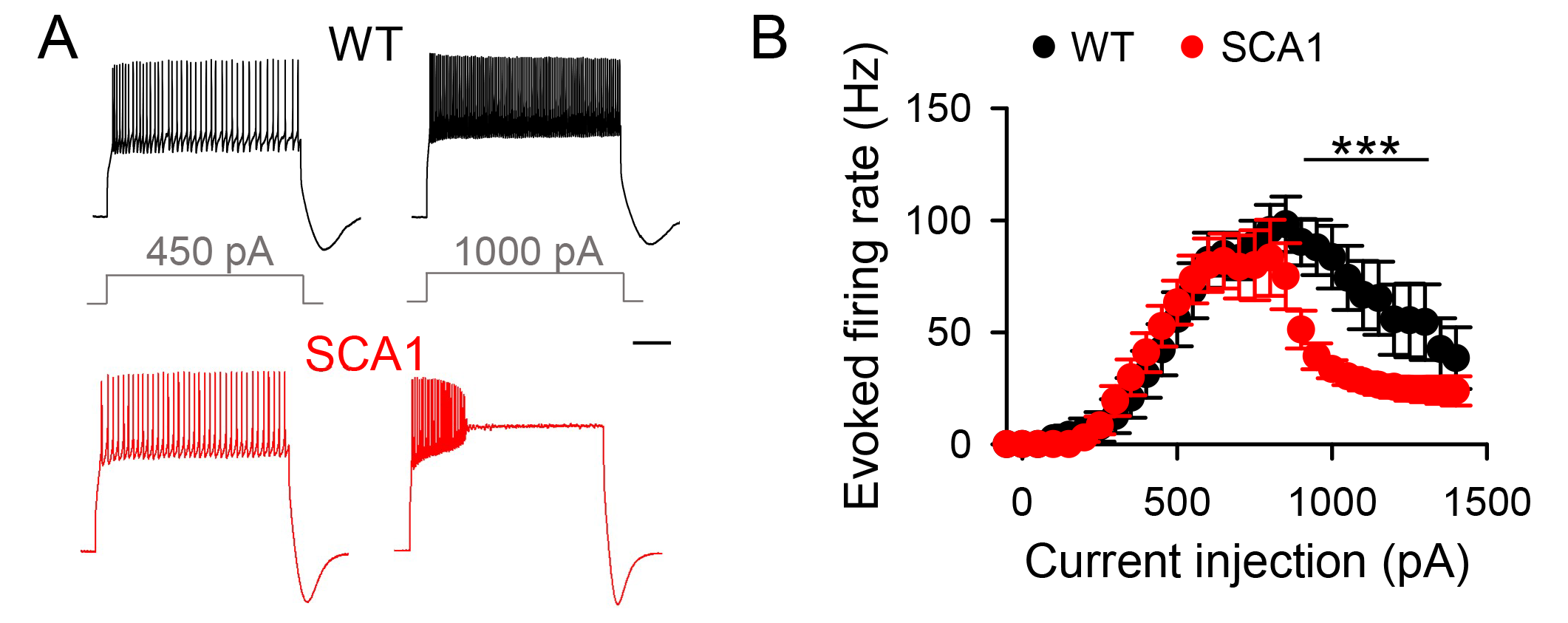
Intrinsic electrical excitability of Purkinje cells is altered in SCA1 mouse model. **A.** Whole-cell recordings of evoked action potentials in PCs from WT (black) and SCA1 (red) mice (scale bar: 200 ms). **B.** Quantitative analysis of evoked PC firing rate (WT, *n* = 17 cells; SCA1, *n* = 13 cells; two-way ANOVA with Tukey post hoc analysis). Note the greater action potential adaptation in SCA1 mice.

#### Afterhyperpolarization currents are altered in SCA1 mouse

We then further investigated the mechanisms underlying the altered intrinsic electrical properties in SCA1 mice. We noticed that PCs in these mice showed a stronger post-burst afterhyperpolarization (AHP) than in WT mice (Figure 4A, B). Three types of AHP currents mediated by calcium-dependent potassium channels have been described, i.e., fast (lasting 2- 5ms), medium (lasting < 200ms) and slow (lasting > 200ms) currents (fI_AHP_, mI_AHP_ and sI_AHP_, respectively) (Storm, 1987; Matthews et al., 2009; Lin et al., 2020). While fI_AHP_ largely contributes to the action potential repolarization (Storm, 1987; Sah, 1996), mI_AHP_ and sI_AHP_ strongly influence neuronal excitability, including spike-frequency adaptation (Andreasen and Lambert, 1995; Sah, 1996; Cohen et al., 1999; Cingolani et al., 2002; Ha and Cheong, 2017). We then asked whether these currents were altered in SCA1 mice. We electrophysiologically recorded PCs in voltage-clamp mode, applied a 800 ms depolarizing pulse to 0 mV from a holding potential of −70 mV and monitored the slowly decaying outward tail current that followed the depolarizing pulse and that contains the mI_AHP_ and sI_AHP_, which were quantified from the peak amplitude of the afterhyperpolarization current and the area under the curve during 1 s after the current peak, respectively (Santini et al., 2008; Maglio et al., 2021). The magnitude of both mI_AHP_ and sI_AHP_ was significantly higher in SCA1 mice than in WT mice (Figure 4C, D). Consistent with the well-known role of mI_AHP_ and sI_AHP_ in regulating neuronal excitability and spike-frequency adaptation, these results suggest that the enhancement of the mI_AHP_ and sI_AHP_ in the SCA1 mice may account for the decreased intrinsic excitability and spontaneous firing activity in PCs of SCA1 mice.

**Figure 4.**
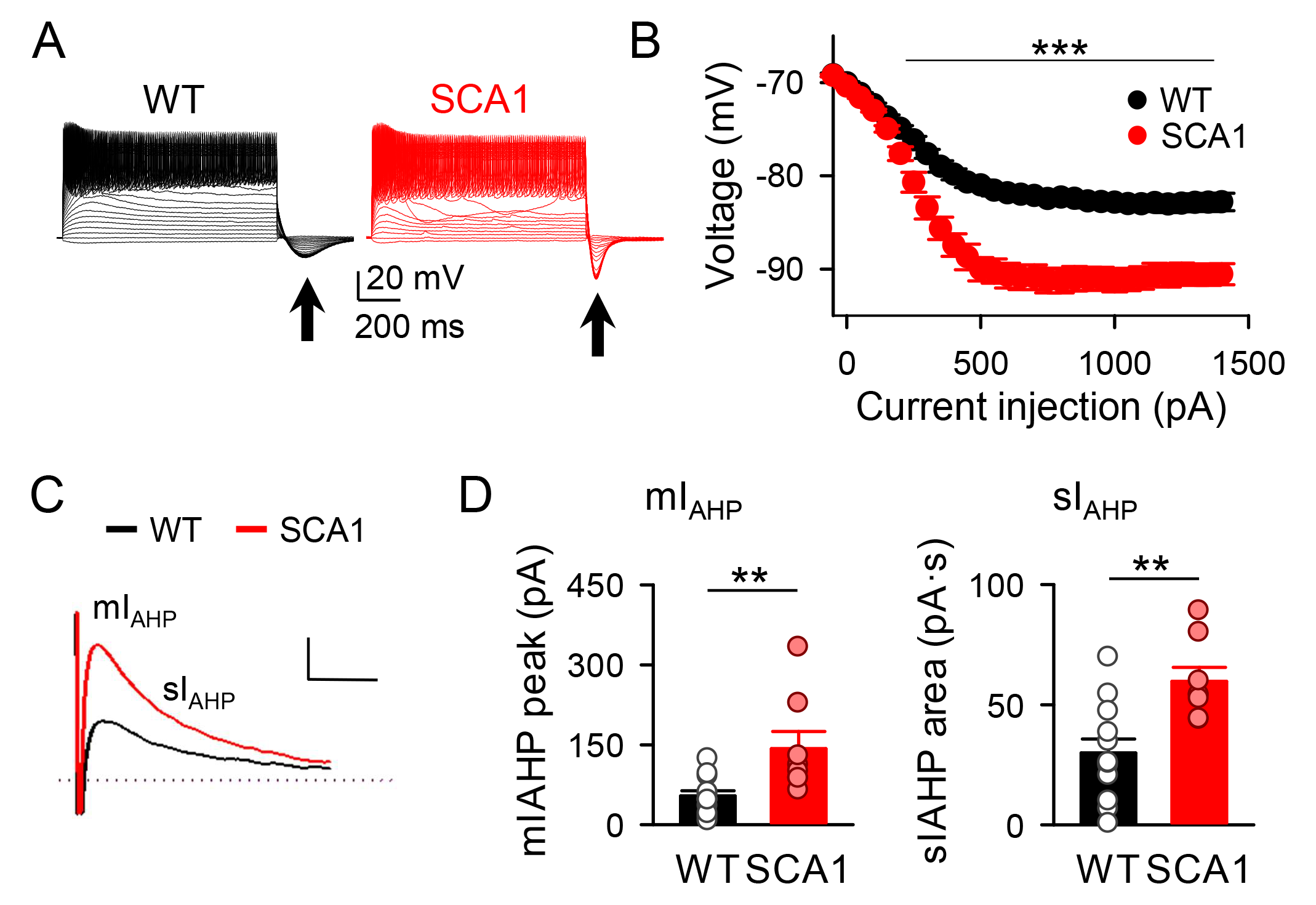
Purkinje cell afterhyperpolarization current is increased in SCA1 mouse model. **A.** Whole-cell recordings of evoked action potentials in PCs from WT (black) and SCA1 (red) mice. Note the afterhyperpolarization current (arrows). **B.** Peak voltage measured with consecutive current injections (WT, *n* = 22 cells; SCA1, *n* = 15 cells; two-way ANOVA with Tukey post hoc analysis). **C.** I_AHP_ representative traces from WT (black) and SCA1 (red) mice (scale bar: 40 pA, 500 ms). **D. Left.** Quantitative analysis of medium I_AHP_ (WT, *n* = 12 cells; SCA1, *n* = 8 cells; Mann-Whitney test; p=0.005). mI_AHP_ is measured as the peak current. **Right.** Quantitative analysis of slow I_AHP_ (WT, *n* = 12 cells; SCA1, *n* = 8 cells; t-test, two-tailed p*=*0.003). sI_AHP_ is measured as the area under the curve during 1 s beginning at the peak of the current.

### Functional properties of Bergmann Glia in SCA1 mice

BG structurally and functionally interact with PCs and synapses in the molecular layer of the cerebellum (Grosche et al., 1999; Brockhaus and Deitmer, 2002; Saab et al., 2012; Wang et al., 2012; Rudolph et al., 2016; Miyazaki et al., 2017). They also enwrap processes of basket and stellate cells (molecular layer interneurons) (Castejón et al., 2002). In addition, BG are equipped with receptors and transporters to sense both inhibitory and excitatory neurotransmitters (De Zeeuw and Hoogland, 2015). BG has been revealed to play important roles in cerebellar function (Saab et al., 2012; Paukert et al., 2014; Salinas□Birt et al., 2023). We therefore asked whether the functional properties of BG were altered in SCA1mice.

#### Spontaneous calcium activity in BG is enhanced in SCA1 mice

BG display intracellular calcium elevations that may occur spontaneously or evoked by neurotransmitters that activate calcium-permeable AMPARs (Iino et al., 2001; Piet and Jahr, 2007), and G-protein-coupled receptors (GPCRs) that stimulate type 2 inositol triphosphate receptor (IP3R2)-dependent calcium mobilization from the endoplasmic reticulum (Araque et al., 2014; Paukert et al., 2014; Rudolph et al., 2016; Kofuji and Araque, 2021; Salinas□Birt et al., 2023). We first analyzed the spontaneous calcium activity of BG in SCA1 and wildtype (WT) mice using the genetically encoded calcium indicator GCaMP6f expressed under the astroglial glutamate transporter (GLAST) promoter. Cerebellar slices were imaged for 5 min, and the spontaneous Ca^2+^ event frequency was analyzed (Figure 5A-C). BG displayed a higher spontaneous Ca^2+^ event frequency in SCA1 mice than in WT mice (0.6 events per min ± 0.1 in WT mice vs 1.1 events per min ± 0.1 in SCA1 mice) (Figure 5B, C). We then tested the BG responsiveness to a puff application (2s) of ATP. Both mice responded to ATP with robust calcium elevations (Figure 5D) and with no significant differences in the ATP-evoked calcium signal, quantified as Ca^2+^ event probability (0.7 ± 0.1 in WT vs 0.7 ± 0.04 in SCA1) (Figure 5E), suggesting that the molecular machinery involved in neurotransmitter-evoked calcium mobilization was unaffected. These results indicate that BG in of SCA1 mice were spontaneously hyperactive.

**Figure 5.**
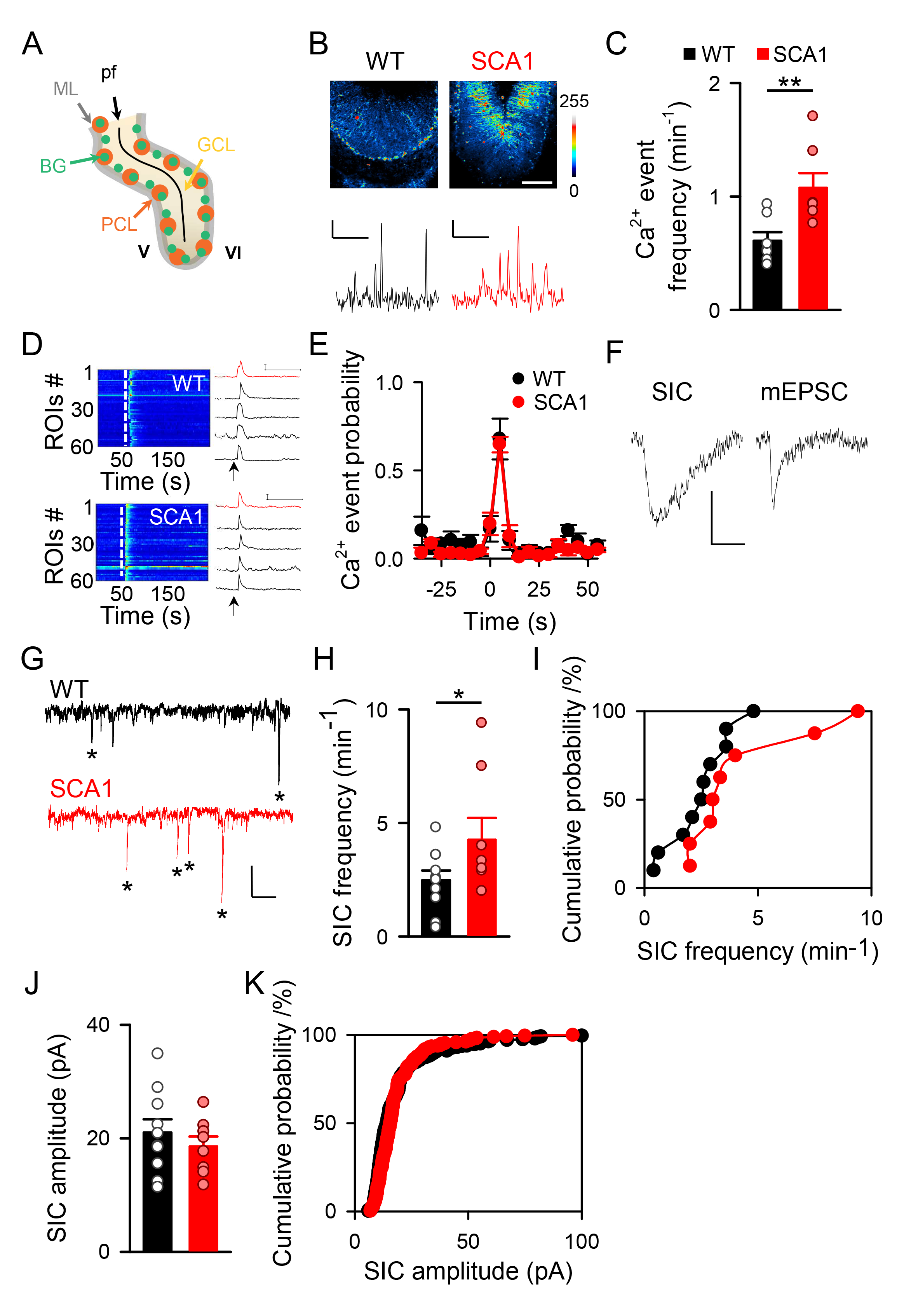
Bergmann glia calcium activity and glutamate release are increased in SCA1 mouse model. **A.** Schematic representation of a sagittal section of the lobules V/VI of the cerebellum of a WT mouse. Lobules are named with roman numbers. Cells were recorded from lobules V and VI at both sides of the primary fissure. pf: primary fissure, ML: Molecular layer, PCL: Purkinje cell layer, GCL: Granule cell layer, BG: Bergmann glia. **B. Top.** Pseudocolor calcium images showing the intensities of GCaMP6 expressing Bergmann glia in WT and SCA1 mice (scale bar: 100 µm). **Bottom.** Representative spontaneous calcium traces from WT (black trace, scale bar: 10%, 100ms) and SCA1 (red trace, scale bar: 5%, 100ms). **C.** Calcium event frequency in WT and SCA1 mice. Note the higher frequency of calcium activity in SCA1 (WT, n = 7 slices; SCA1, n = 7 slices; Mann-Whitney test; p=0.007). **D. Left.** Representative heat map plots of calcium events in response to an ATP puff (white dash line) in WT and SCA1. **Right.** Calcium traces in response to an ATP puff (arrows) and the average (red trace) in WT (Top, scale bar: 20%, 100 s) and SCA1 (bottom, scale bar: 40%, 100 s). **E.** Calcium event probability during a puff of ATP. (WT, n = 4 slices; SCA1, n = 4 slices; t-test at bin 5s, two-tailed p=0.81). **F.** Representative trace of a SIC and a miniature EPSC (scale bar: 10 pA, 25 ms). Note the difference in the kinetics, with the SIC being slower than the mEPSC. **G.** Representative traces of slow inward currents (SICs, asterisks) recorded from a Purkinje cell (PC) from a WT (black) and SCA1 (red) mouse (scale bar: 10 pA, 2 s). **H.** SIC frequency in WT and SCA1 mice. Note the higher frequency of SICs in SCA1 (WT, n = 10; SCA1, n = 8; t-test, one-tailed p=0.043). **I.** Cumulative probability of SIC frequency from the cells recorded in H. **J.** SIC amplitude in WT and SCA1 mice (*WT*, n = 10; *SCA1*, n = 8; t-test, two-tailed p=0.44). **K.** Cumulative probability of SIC amplitude from the cells recorded in H.

#### Glutamatergic gliotransmission is enhanced in SCA1 mice

Astroglial calcium elevations are known to stimulate the release of gliotransmitters in different brain areas (Volterra and Meldolesi, 2005). In the cerebellum, activation of BG induces the release of the gliotransmitter glutamate (Rudolph et al., 2016; Shim et al., 2018; Beppu et al., 2021). We then investigated whether the BG calcium hyperexcitability in SCA1 mice was accompanied by an enhanced glutamatergic gliotransmission, monitoring the calcium-dependent NMDAR-mediated slow inward currents (SICs) in adjacent neurons, as a biological assay of glutamate release (Araque et al., 1998; Angulo, 2004; Fellin et al., 2004; Perea and Araque, 2005; Navarrete and Araque, 2008). We recorded whole-cell currents from PCs in the presence of TTX (1 µM) and in the absence of extracellular Mg^2+^ to maximize NMDAR activation. In agreement with previous reports (Araque et al., 1998; Angulo, 2004; Fellin et al., 2004; Perea and Araque, 2005; Navarrete and Araque, 2008), SICs could be differentiated from spontaneous and miniature excitatory postsynaptic currents (sEPSCs and mEPSCs, respectively) by their significant slower kinetics (Figure 5F). SICs were abolished in the presence of AP5 (50 µM), confirming that they were mediated by NMDAR activation (Angulo, 2004; Fellin et al., 2004; Perea and Araque, 2005) (data not shown). The frequency of SICs was significantly higher in SCA1 than in WT mice (4.3 ± 1.0 and 2.5 ± 0.4 min^-1^, respectively) (Figure 5G-I), whereas no statistical differences were found in the SIC amplitude (21.0 ± 2.3 pA in WT vs 18.6 ± 1.7 pA in SCA1 mice) (Figure 5J, K). Altogether, these results indicate an enhancement of glutamatergic gliotransmission, which is consistent with the calcium hyperexcitability of BG, in SCA1 mice.

#### Glutamate uptake and K^+^ buffering by BG is not altered SCA1 mice

The uptake of neurotransmitters (Anderson and Swanson, 2000; Schousboe, 2000), and the buffering of extracellular potassium (Sofroniew and Vinters, 2010; Clarke and Barres, 2013; Khakh and Sofroniew, 2015; Farhy-Tselnicker and Allen, 2018; Magaki et al., 2018) are two crucial roles of astroglia. Hence, we tested whether these functions were altered in BG of SCA1 mice.

First, we monitored the synaptically-evoked glutamate transporter currents in visually and electrophysiologically identified BG (see Methods). We whole-cell electrophysiologically recorded BG in voltage-clamp conditions and stimulated nearby synapses with an extracellular bipolar electrode using trains (500 ms) of stimuli (at 75 Hz) at different intensities (Figure 6A). Because glutamate transporter activity is electrogenic (Kanai et al., 1995; Zerangue and Kavanaugh, 1996; Grewer and Rauen, 2005), the stimulation evoked slow currents that were largely mediated by glutamate transporter activation, as confirmed by their sensitivity to Threo-β-benzyl-oxyaspartate

**Figure 6.**
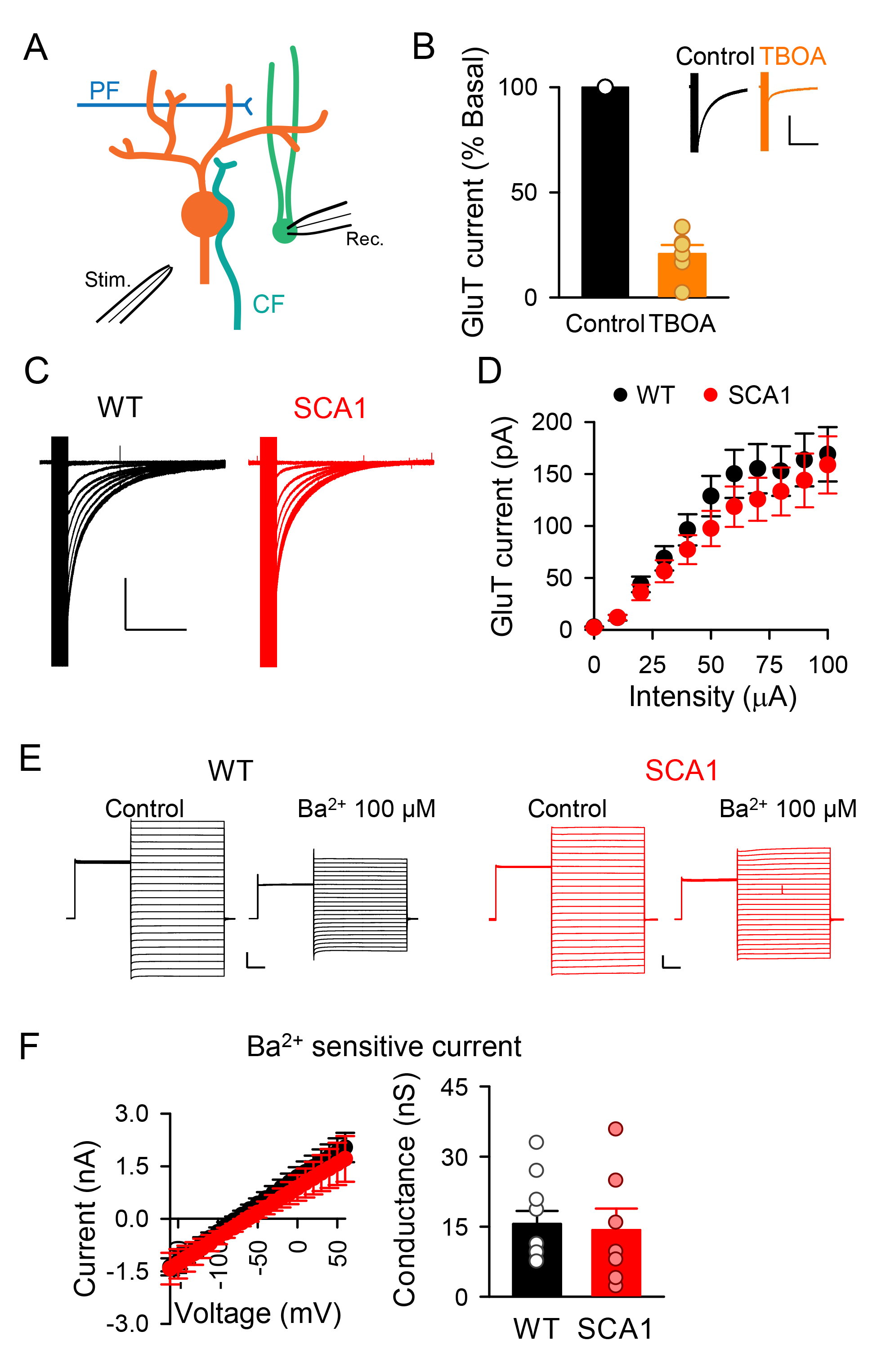
Bergmann glia glutamate transport current and Ba^2+^ sensitive currents are not altered in SCA1 mouse model. **A.** Scheme of the experimental approach. Glutamatergic climbing (turquoise) and parallel (blue) terminals are stimulated with a bipolar stimulator (Stim.). Glutamate transporter currents are recorded from Bergmann glia (green). Purkinje cell is depicted in orange. **B.** Glutamate transporter current during basal conditions and in the presence of TBOA (10 µM, *Control*, n = 6; TBOA, n = 6; paired t-test; p<0.001) (scale bar: 100 pA, 2 s). **C.** Representative traces of the glutamate transporter current evoked at different intensities of stimulation in WT and SCA1 mice (scale bar: 100 pA, 2 s). **D.** Input-Output relationships of glutamate transporter current evoked by different intensities of stimulation (*WT*, n = 11; *SCA1*, n = 16; two-way ANOVA with Tukey *post hoc* analysis). **E.** Representative traces obtained at different voltages steps before and after barium application in Bergmann glia from WT and SCA1 mice (scale bar: 1 nA, 100 ms). Note the blockage of the current produced by barium perfusion. **F. Left.** Barium-sensitive current in WT and SCA1 mice (*WT*, n = 10; *SCA1*, n = 7; two-way ANOVA with Tukey *post hoc* analysis). **Right.** Conductance (measured in picosiemens(pS)) of the barium-sensitive current in WT and SCA1 mice (*WT*, n = 10; *SCA1*, n = 7; Mann-Whitney test; p=0.591).

(TBOA; 10 µM) (Shimamoto et al., 1998) (Figure 6B). The amplitude of the synaptically-evoked glutamate transporter currents in WT and SCA1 mice were not significantly different (Figure 6C, D), indicating that glutamate uptake function of BG was not compromised in SCA1 mice.

Then, we assessed the potassium buffer capacity of BG by monitoring the functional expression of Kir4.1 channels, Ba^2+^-sensitive inwardly rectifying potassium channels that are largely expressed in astroglia and that are critical in maintaining the extracellular K^+^ concentration. We electrophysiologically recoded BG in voltage-clamp conditions and applied the following protocol to monitor the whole-cell currents maximizing the inwardly rectifying currents: a 300 ms pre-pulse to 0 mV, followed by 500 ms pulses of increasing voltages (from −160 to +60 mV in 10 mV steps). BaCl_2_ (100 µM) was then perfused to isolate the Ba^2+^-sensitive Kir4.1 currents by subtraction of the currents. No differences were found in the I-V curves of the Ba^2+^ sensitive currents and their slopes between WT and SCA1 mice (Figure 6E, F), indicating that the functional expression of Kir4.1 channels, suggesting that the potassium buffer capacity of BG was unaltered in SCA1 mice.

Altogether, these results indicate that the crucial supportive roles of BG, the glutamate uptake and ion homeostasis, were not compromised in SCA1 mice.

#### Calcium signal in BG regulates spontaneous firing rate and PC excitability

Because BG functionally interact with PCs and their synaptic inputs (Grosche et al., 1999; Brockhaus and Deitmer, 2002; Saab et al., 2012; Wang et al., 2012; Rudolph et al., 2016; Miyazaki et al., 2017), we hypothesized that the BG calcium hyperactivity may contribute to the alterations of the PC electrical intrinsic properties and, consequently, the spontaneous firing rate in SCA1 mice. To test this hypothesis, we first recorded the mI_AHP_ and sI_AHP_ in a PC before and after loading a nearby BG (<120 µm; see Methods for details) with the Ca^2+^ chelator BAPTA through a patch-clamp recording pipette (Figure 7A, B). BAPTA loading in BG by did not affect mI_AHP_ or sI_AHP_ in WT mice (mI_AHP_: 45.3±13.6 pA and 43.3±4.9 pA; sI_AHP_: 22.6±7.2 pA and 20.7±2.9 pA, in control and BAPTA-loaded BG, respectively) (Figure 7C, D). In contrast, in SCA1 mice, the enhanced mI_AHP_ and sI_AHP_ were reduced by BG BAPTA loading, attaining similar values than WT mice (mI_AHP_: 105.67±6.91 pA and 50.61±13.81 pA; sI_AHP_: 55.67±5.81 pA vs 17.82±5.66 pA, in control and BAPTA-loaded BG, respectively (Figure 7C, D). These results indicate that preventing the Ca^2+^ activity in BG in SCA1 mice reduced the enhanced mI_AHP_ or sI_AHP_, restoring their values to those of WT mice.

**Figure 7.**
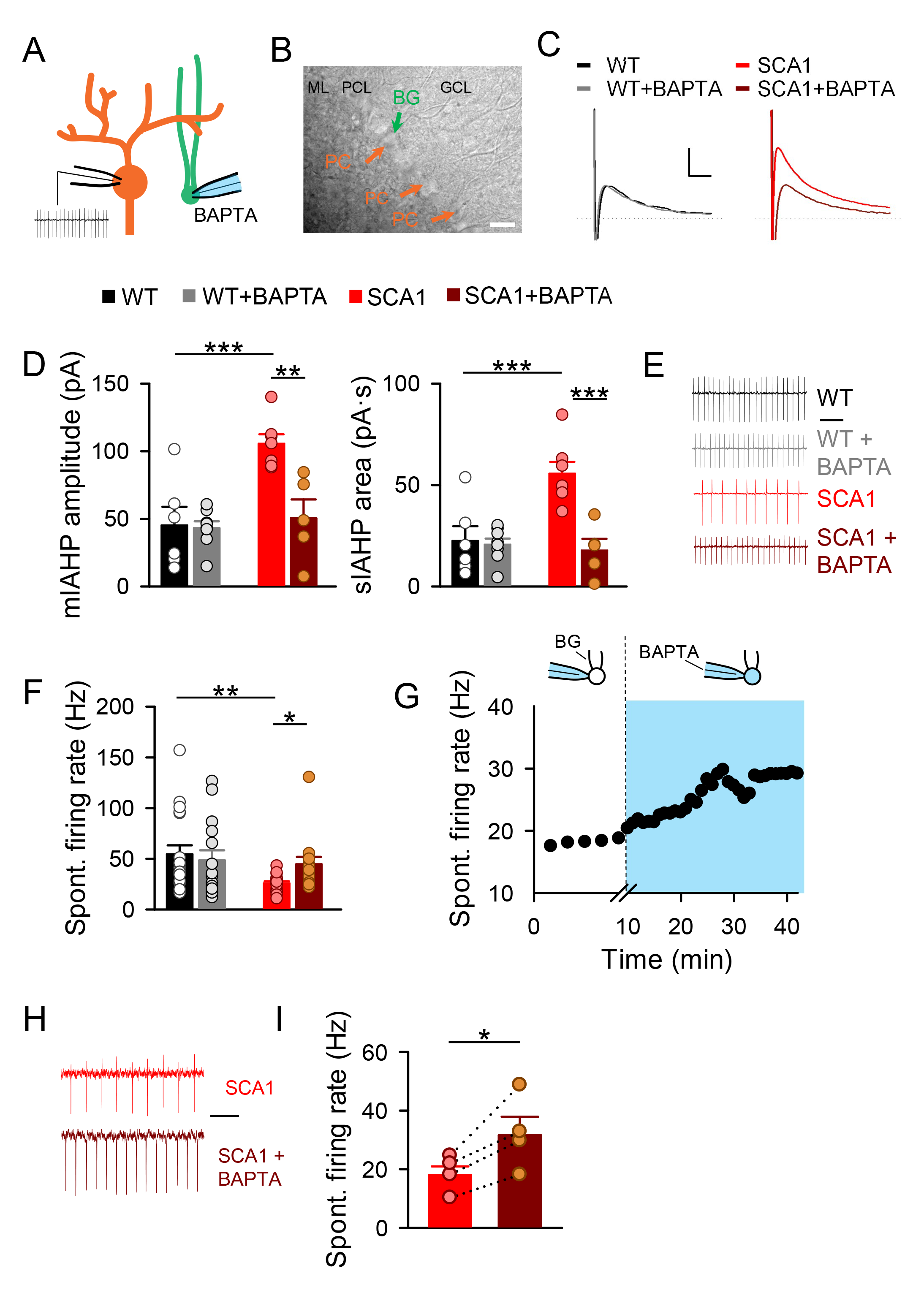
Calcium chelation in Bergmann glia restores alterations found in SCA1 mouse model. **A.** Scheme of the experimental approach. A BG (green) is loaded with BAPTA through the patch pipette. After allowing the BAPTA to diffuse different parameters are measured in a nearby PC (orange). **B.** Differential interference contrast (DIC) image of a sagittal cerebellar section of a WT mouse. A BG is loaded with BAPTA (green arrow) and allowed to diffuse through the BG network. Recorded PCs (orange arrows) were no more than 120 µm away from the loaded BG (scale bar: 25 µm). **C.** Representative traces of mI_AHP_ and sI_AHP_ in PCs from WT (black), WT with BAPTA-loaded BG (grey), SCA1 (red), and SCA1 with BAPTA-loaded BG (dark red) mice (scale bar: 40 pA, 500 ms). **D. Left.** Quantitative analysis of medium I_AHP_ (WT, *n* = 6 cells; WT+BAPTA, *n* = 8 cells; SCA1, *n* = 7 cells; SCA1+BAPTA, *n* = 5 cells; one-way ANOVA with Tukey *post hoc* analysis). mI_AHP_ is measured as the peak current. **Right.** Quantitative analysis of slow I_AHP_ (WT, *n* = 6 cells; WT+BAPTA, *n* = 8 cells; SCA1, *n* = 7 cells; SCA1+BAPTA, *n* = 5 cells; one-way ANOVA with Tukey *post hoc* analysis). sI_AHP_ is measured as the area under the curve during 1 s beginning at the peak of the current. **E.** Cell-attached recordings of spontaneous action potential firing in PCs from WT (black), WT with BAPTA-loaded BG (grey), SCA1 (red), and SCA1 with BAPTA-loaded BG (dark red) mice (scale bar: 100 ms). **F.** Quantitative analysis of PC firing rate in the mentioned different conditions (WT, *n* = 19 cells; WT+BAPTA, *n* = 15 cells; SCA1, *n* = 19 cells; SCA1+BAPTA, *n* = 14 cells; Kruskal-Wallis one-way ANOVA with Dunńs *post hoc* analysis; WT vs SCA1 p=0.007; SCA1 vs SCA1+BAPTA p=0.033). **G.** A BG and a nearby PC were patched at the same time. After recording the spontaneous firing rate for a baseline period of 5 min in the PC, the seal in the BG was ruptured and the BAPTA inside the pipette was allowed to diffuse (shaded in blue) while recording the spontaneous firing rate in the PC. Note that the firing rate in the SCA1 PC increases as the BAPTA is diffusing into the BG. An omission in the x axis (Time) from minute 5 to 10 was made to better appreciate the baseline period. **H.** Cell-attached recordings of spontaneous action potential firing rate from the PC recorded in G before (red) and after the BAPTA diffusion (dark red) (scale bar: 100 ms). **I.** Quantitative analysis of PC firing rate in SCA1 mice before (red) and after (dark red) BAPTA diffusion (SCA1, *n* = 4 cells; SCA1+BAPTA, *n* = 4 cells; paired t-test, two-tailed *P =* 0.032).

Because mI_AHP_ and sI_AHP_ strongly influence neuronal excitability and spike-frequency adaptation (Ha and Cheong, 2017), we then investigated whether BAPTA loading of BG affected the spontaneous firing rate in SCA1 mice. Consistent with previous results (see Fig. 1A, B), in control conditions, the spontaneous firing rate of PC in WT mice was significantly higher than in SCA1 mice (Figure 7E, F). After BAPTA loading (for at least 30 min) of an adjacent BG (distance of the somas: 50±6 µm), the spontaneous firing rate was unaffected in WT mice (54.8±8.9 Hz in control and 48.7±10.0 Hz after at least 30 min of BAPTA-loaded BG), but it was significantly increased in SCA1 mice (26.0±1.9 Hz in control and 44.8±7.4 Hz in BAPTA-loaded BG) to attain values close to WT mice (Figure 7E, F). To discard potential sampling effects due to the variability of the spontaneous firing rate among PCs, we aimed to confirm these results by performing paired-recordings of the PC and the BG (both in cell-attached patch-clamp mode) to continuously monitor the spontaneous firing activity in control conditions (baseline period of 5 min) and after breaking the gigaohm seal in the BG that allowed the intracellular diffusion of BAPTA in the BG in SCA1 mice (Figure 7F). In agreement with the above results, the spontaneous firing rate in the PC was increased after loading the BG with BAPTA (from 18.0 ± 3.4 Hz to 31.6 ±7.3 Hz after 30 min of BAPTA loading; n = 4) (Figure 7F, G, H and I). These results indicate that preventing the BG calcium activity in SCA1 mice increased the spontaneous firing rate of PC, suggesting that the BG hyperactivity contributes to the impaired PC firing activity in SCA1 mice.

It is known that increase calcium in astrocytes induces glutamate-mediated slow inward currents (SICs) in adjacent neurons (Araque et al., 1998; Angulo, 2004; Fellin et al., 2004; Perea and Araque, 2005; Navarrete and Araque, 2008). Thus, preventing calcium elevation in the BG should decrease the SIC frequency. Accordingly, we found a decrease in the SIC frequency in both WT and SCA1 mice after loading BG with BAPTA (Supplementary Figure 1A, B). This effect was more prominent in SCA1 mice. SIC frequency in WT mice was reduced from 2.0±0.6 min^-1^ to 1.4±0.3 min^-1^, while in SCA1 mice SIC frequency varied from 3.9±0.6 min^-1^ to 1.0±0.3 min^-1^ after loading BG with BAPTA (Supplementary Figure 1A, B).

#### Exogenous activation of BG in WT recapitulates the impairments of PCs in SCA1 mice

Previous results indicate that the calcium hyperactivity of BG contributes to the enhancement of mIAHP and sIAHP and the consequent decrease of the firing activity of PCs in SCA1 mice. To further test this idea, we asked whether exogenous activation of the BG using Designer Receptors Exclusively Activated by Designer Drugs (DREADDs) in WT mice would mimic the alterations observed in SCA1 mice. For that purpose, WT mice were injected with AAV8-GFAP-Gq-DREADD-mCherry in lobule VI of the cerebellum (Figure 8A, B). We then tested whether these calcium elevations stimulated gliotransmission by recording the glutamate-mediated SICs in PCs 10 min before and after perfusing CNO. In WT mice, CNO significantly increased SIC frequency (from 2.3±0.3 min^-1^ to 4.4±0.5 min^-1^) (Figure 8C, D). In contrast, in SCA1 mice, the already high SIC frequency (4.4±1.3 min^-1^) was not altered by CNO (4.2±0.6 min^-1^) (Supplementary Figure 1C, D), suggesting a saturated process due to the BG calcium hyperactivity in these mice (see Fig. 5C). We then investigated whether mIAHP and SIAHP were affected by activating DREADD-expressing BG with CNO. In WT mice, both currents were increased after CNO perfusion (mI_AHP_: 42.4±9.3 pA and 89.8±17.4 pA, before and after CNO perfusion, respectively; sI_AHP_: 19.4±4.2 pA·s and 41.9±9.4 pA·s before and after CNO perfusion, respectively) (Figure 8E, F). In contrast, as expected form the saturating calcium hyperactivity, both currents were unaffected by CNO in SCA1 mice (mI_AHP_: 103.5±10.7 pA and 129.8±18.1 pA, before and after CNO perfusion, respectively; sI_AHP_: 58.9±6.0 pA·s vs 69.8±9.7 pA·s, before and after CNO perfusion, respectively) (Supplementary Figure 1E, F). These results indicate that mIAHP and sIAHP were regulated by activation of the BG calcium signal in WT mice, attaining similar high values as in SCA1 mice.

**Figure 8.**
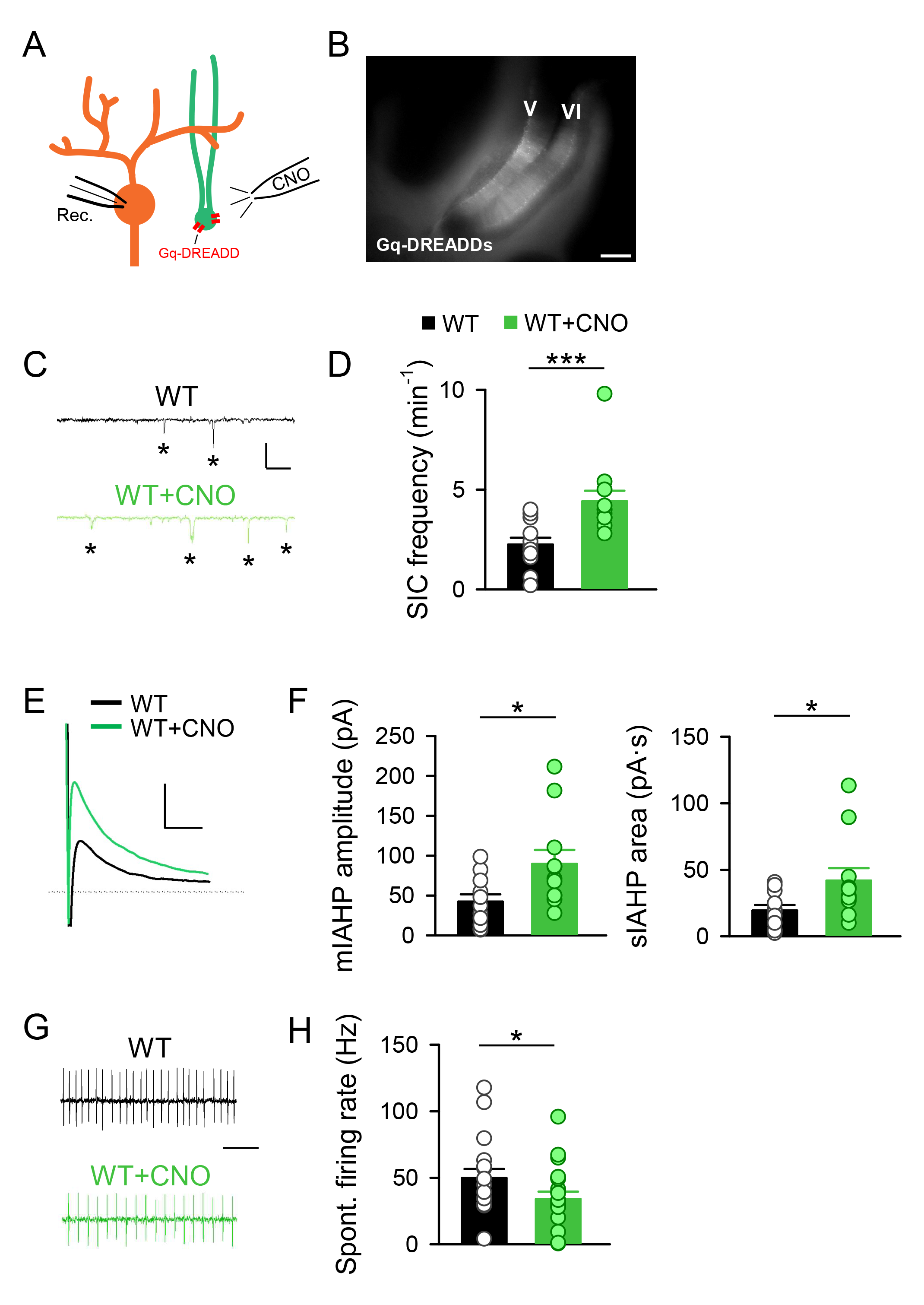
DREADDs activation in Bergmann glia from WT mice mimic the alterations found in SCA1 mice. **A.** Scheme of the experimental approach. Gq-DREADDs are expressed in Bergmann glia (green) and activated via the perfusion of CNO. The effect of BG activation is measured in nearby PCs (orange). **B.** GFAP-Gq-DREADD-mCherry expression after viral injection in the cerebellum. Note that the virus is expressed in lobules V and VI (scale bar: 250 µm). **C**. Representative traces of SICs in PCs from WT (black) and WT+CNO (green) mice (scale bar: 40 pA, 2 s). **D.** Quantitative analysis of SIC frequency (WT, *n* = 12 cells; WT+CNO, *n* = 12 cells; Mann-Whitney test; p=0.001). **E.** Representative traces of mI_AHP_ and sI_AHP_ in PCs from WT (black) and WT+CNO (green) mice (scale bar: 40 pA, 500 ms). **F. Left.** Quantitative analysis of medium I_AHP_ (WT, *n* = 11 cells; WT+CNO, *n* = 11 cells; Mann-Whitney test; p=0.026). mI_AHP_ is measured as the peak current. **Right.** Quantitative analysis of slow I_AHP_ (WT, *n* = 11 cells; WT+CNO, *n* = 11 cells; Mann-Whitney test; p=0.042). sI_AHP_ is measured as the area under the curve during 1 s beginning at the peak of the current. **G.** Cell-attached recordings of spontaneous action potential firing in PCs from WT (black), WT+CNO (green) mice (scale bar: 100 ms). **H.** Quantitative analysis of PC spontaneous firing rate in the mentioned different conditions (WT, *n* = 18 cells; WT+CNO, *n* = 20 cells; t-test, one-tailed p=0.038).

We finally tested whether PC firing rate was affected by stimulating the calcium activity in BG-expressing DREADDs with CNO. In WT mice, BG stimulation with CNO decreased the PC firing rate (50.0±6.9 Hz and 34.1±5.8 Hz, before and after CNO, respectively) (Figure 8G, H). Contrastingly, and consistently with the saturating calcium hyperactivity, in SCA1 mice, CNO did not affect the already decreased PC firing rate (21.0±2.2 Hz and 25.6±2.5 Hz, before and after CNO, respectively) (Supplementary Figure 1G, H), indicating that BG stimulation in WT mice mimicked the reduced firing rate in SCA1 mice.

Taken together, these results indicate that selective BG activation reproduced the cellular phenotype of SCA1 mice, i.e., BG calcium activation, up-regulation of mIAHP and sIAHP function, and decrease PC firing rate.

## Discussion

In the present study, we combined electrophysiological, calcium imaging and chemogenetic approaches in cerebellar brain slices to investigate the cellular phenotypic alterations of PCs and BG in a mouse model of spinocerebellar ataxia type 1 (SCA1). We report here that SCA1 PCs showed a lower spontaneous firing rate and larger medium and slow afterhyperpolarization currents (mI_AHP_ and sI_AHP_) than wildtype mice. SCA1 BG displayed a dysfunctional calcium hyperactivity and gliotransmission, manifested by a higher frequency of NMDAR-mediated slow inward currents (SICs) in PC that can be reverted by manipulating the calcium activity in BG. Preventing BG hyperexcitability in SCA1 mice BG regulates the intrinsic properties of PC in the SCA1 mouse model. Besides, mimicking the BG hyperactivity by activating BG expressing Gq-DREADDs in wildtype mice (WT) enhanced mI_AHP_ and sI_AHP_ and the frequency of SICs, as well as reduced spontaneous firing rate, reproducing the SCA1 pathological phenotype of PCs. The role of firing rate coding of the PCs in the cerebellum is well known. The spontaneous firing rate of PCs is regulated by excitatory and inhibitory synaptic inputs for the integration and propagation of the information (Armstrong and Rawson, 1979; Häusser and Clark, 1997). The loss of PCs arborization in SCA1 mouse model compromises the neuronal integrity to exert an effective neuronal function, leading to a progressive dysfunctional firing rate (Inoue et al., 2001; Barnes et al., 2011; Hourez et al., 2011; Dell’Orco et al., 2015). In this study, we show a reduction in the spontaneous firing rate of SCA1 PCs compared with WT mice (Figure 1C-E) as well as the inability of SCA1 PCs to sustain long trains of action potentials in response to depolarizing pulses (Figure 3). The firing rate relies on a balance between depolarizing sodium channels and repolarizing potassium channels. Calcium-activated potassium channels mediating afterhyperpolarization currents (I_AHP_) play a key role in the regulation of firing rate. Reduction in BK channels-mediated small I_AHP_ has being described as an underlying mechanism in the dysfunctional intrinsic excitability of SCA1 PCs (Dell’Orco et al., 2015). Here we reveal a higher magnitude in the medium and slow I_AHP_ (mI_AHP_ and sI_AHP_) of SCA1 PCs than wildtype mice (Figure 4C, D). The enhancement of mI_AHP_ and sI_AHP_ in SCA1 PCs explains the lower firing rate and, therefore, compromising the ability of PCs to regulate the intrinsic properties and information processing.

Cellular systematization of the cerebellum makes it an organized structure. The whole input reaching the cerebellum converges on the PCs receiving afferents either to their somas from climbing fibers or to their dendritic arbors from parallel fibers (Napper and Harvey, 1988; Voogd and Glickstein, 1998; Wu et al., 1999; Sugihara, 2005) (Figure 2A). The study of the synaptic transmission onto the PCs revealed a rather conserved circuitry in the cerebellum of SCA1 mice (Figure 2). Our results show that excitatory synaptic transmission coming from both the climbing and parallel fibers was not altered in the ataxic mouse model. This is in line with the previous studies which displayed no major abnormalities in parallel or climbing fibers to PC innervation (Inoue et al. 2001; Watase et al. 2002; Barnes et al. 2011; Shuvaev et al., 2016). The results here described suggest that basal AMPA receptor-mediated fast synaptic transmission to PCs remains nearly intact in SCA1 mice. In discrepancy with our results, there are some studies showing impairments in the excitatory transmission onto PCs (For a review see Hoxha et al., 2018) (but see also Hourez et al. 2011). This is likely due to the use of different mouse models in which the mutant Ataxin-1 is expressed only in PCs or through the brain. Additionally, ataxia is a very broad disease, which may be caused by different mechanisms or a sum of mechanisms. Thus, alterations in different cell types on the circuitry could lead to similar symptoms.

Bergmann glia is responsible for glutamate uptake and extracellular K^+^ homeostatic clearance (Bellamy, 2006). Astrocytic Glutamate transporter-1 (GLT-1) and inwardly rectifying potassium channels (Kir) play a major role in glutamate removal from the synapses and spatial buffering of K^+^ released by neurons during action potential propagation (Orkand et al., 1966; Newman et al., 1984; Newman, 1986, 1993; Kofuji and Newman, 2004; Wang et al., 2012). In spite of a plethora subtypes of Kir in glial cells, the evidence suggests that Kir4.1 is the principal pore forming subunit in glial cells (Neusch et al., 2001, 2006; Olsen et al., 2006; Djukic et al., 2007). Some evidence shows a downregulation of Kir and GLT-1 expression in SCA1 mouse model (Cvetanovic, 2015; Dell’Orco et al., 2015; Shuvaev et al., 2021 Borgenheimer et al 2022). Similarly, a reduction in the glutamate transporter function has been shown when studying cultured BG and cerebellar synaptosomes derived from different mice models. However, this reduction was not observed using cerebellar slices (Custer et al., 2006). We did not observe differences in the GLT-1 and the Ba^2+^ sensitive K^+^ currents activity between WT and SCA1 mice (Figure 6). Despite the discrepancies, the more likely explanations are the use of brain slices described in previous studies (Custer et al., 2006).

It is extensively established that astrocyte calcium signaling in many brain regions modulates synaptic activity. Astrocyte reactivity in response to a pathological condition, such as an injury or disease, leads to morphological and functional changes. (Takano et al., 2007; Kuchibhotla et al., 2009; Delekate et al., 2014; Angelova et al., 2016; Jiang et al., 2016; Booth et al., 2017; Gómez-Gonzalo et al., 2017; Lines et al., 2022; Nanclares et al., 2023). Early BG reactivity has been described in SCA1 mouse models, contributing to disease pathogenesis (Qu et al., 2017; Kim et al., 2018). Our results show BG calcium hyperactivity exhibited as an increase in spontaneous Ca^2+^ frequency in SCA1 (Figure 5 A-C,). Although astroglial activation have been reported in ataxia models (Seki et al., 2018; Sheeler et al., 2021), this is the first study showing an increase in the calcium signaling within the BG. Additionally, calcium increases in astroglia induce glutamate-mediated slow inward currents (SIC) in adjacent neurons (Araque et al., 1998; Angulo, 2004; Fellin et al., 2004; Perea and Araque, 2005; Navarrete and Araque, 2008). We have found dysfunctional BG gliotransmission, manifested as higher frequency of SICs in cerebellar slices of SCA1 PCs compared with wildtype mice (Figure 4F-K). SCA1 BG hyperexcitability and dysfunctional gliotransmission were reverted manipulating intracellular Ca^2+^ signaling by loading BG with the Ca^2+^ chelator BAPTA, as well as spontaneous firing rate (Supplementary Figure 1, Figure 7E-I), and mI_AHP_ and sI_AHP_ (Figure 7C, D) of PCs. This is one of the first demonstrations of the regulation of neuronal intrinsic properties by glial cells. Strikingly, chemogenetic activation of WT BG expressing Gq-DREADDs induced an increase in the SIC frequency measured in PCs (Figure 8C, D). It also led to a decrease in the spontaneous firing rate on PCs (Figure 8G, H) and showed an increase in both mI_AHP_ and sI_AHP_ in WT PCs (Figure 8E, F), reproducing the SCA1 pathological phenotype of PCs.

While physiological function of ATXN1 is not completely understood, polyQ expansion in ataxin-1 causes perturbations to intracellular homeostatic processes (Irwin et al., 2005) leading to a SCA1. This is characterized by progressive motor deficits, cognitive decline, and mood changes and it has been associated with a severe degeneration of cerebellar PCs and cerebellar gliosis of BG. Here, we show that PCs from SCA1 mice display a lower spontaneous firing rate than WT supported by the higher magnitude of mI_AHP_ and sI_AHP._ In addition, hyperexcitability of BG calcium activity in SCA1 mice together with the dysfunctional gliotransmission show a non-cell type specific altered mechanism in the progression of SCA1. The manipulation of BG activity through intracellular Ca^2+^ BAPTA reverted the intrinsic properties of PC to values of WT mice, as well as restored gliotransmission, demonstrating that BG regulates neuronal intrinsic properties. Interestingly, the activation of BG expressing Gq-DREADDs in WT mice mimicked the pathological phenotype in PC. Despite further studies are necessary, our results reveal a novel dysfunctional mechanism of BG-PC network in SCA1 mice.

## Conclusion

SCA1 BG dysfunction influences the alteration of the intrinsic properties of SCA1 PCs. This malfunction was prevented by manipulating SCA1 BG hyperactivity, reversing the functionality of the intrinsic properties of PCs to values similar to those of WTs. Interestingly, chemogenetic activation of BG mimicked the pathological phenotype in WT PCs. These findings are one of the first pieces of evidence on the regulation of the intrinsic properties of neurons by glial cells. Therefore, Bergmann glia has a relevant function in the control of intrinsic properties of PC in SCA 1 and may serve as potential targets for therapeutic approaches to treat this disease.

---

**Supplementary Figure 1.**
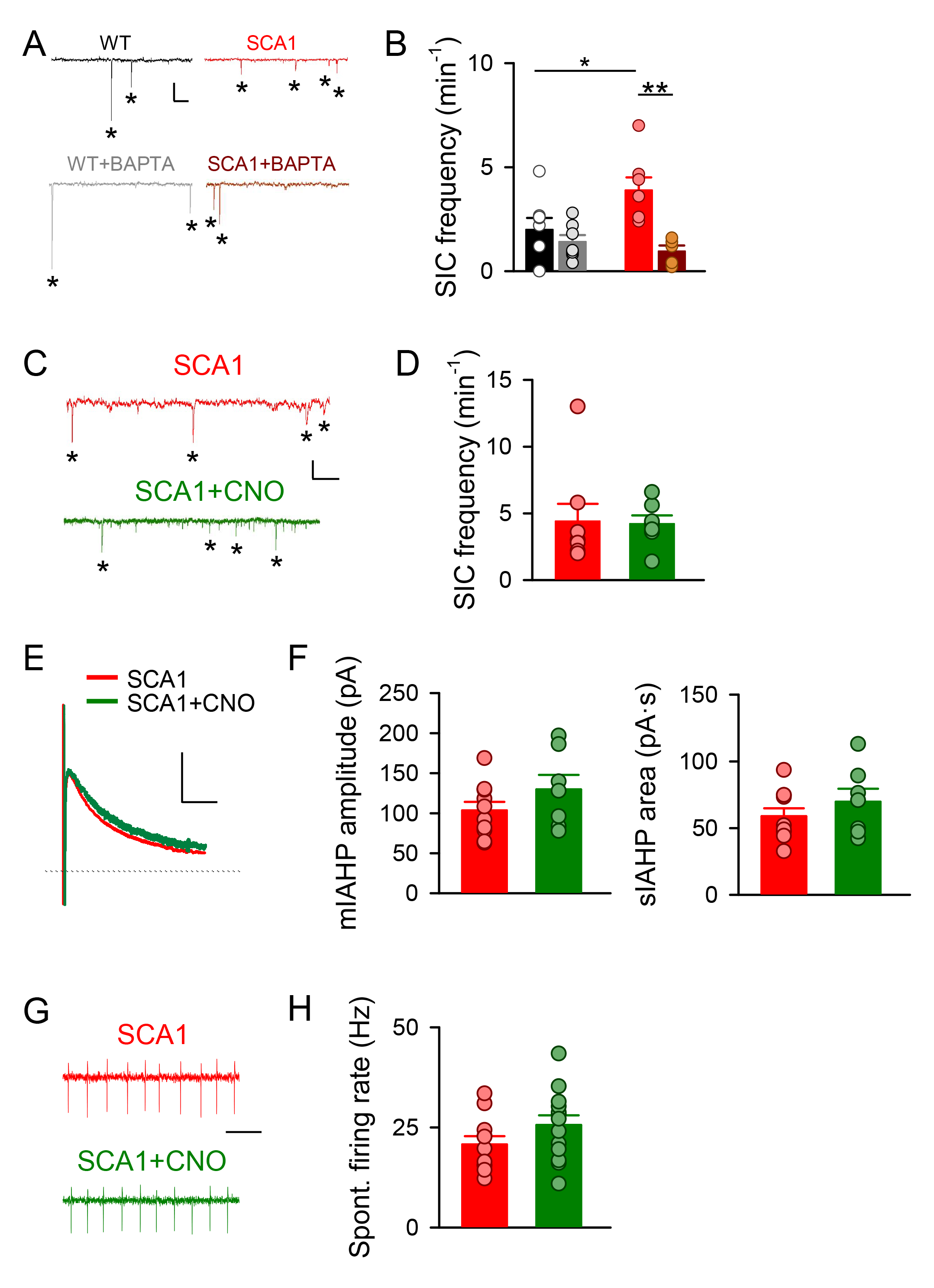
Calcium chelation in Bergmann glia reduces their glutamate release while DREADDs activation in Bergmann glia from SCA1 mice has no effect. **A.** Representative traces of slow inward currents (SICs) in PCs from WT (black), WT with BAPTA loaded BG (grey), SCA1 (red), and SCA1 with BAPTA loaded BG (dark red) mice (scale bar: 40 pA, 2 s). **B.** Quantitative analysis of SIC frequency (WT, *n* = 8 cells; WT+BAPTA, *n* = 7 cells; SCA1, *n* = 7 cells; SCA1+BAPTA, *n* = 5 cells; one-way ANOVA with Tukey *post hoc* analysis). **C.** Representative traces of SICs in PCs from SCA1 (red) and SCA1+CNO (dark green) mice (scale bar: 40 pA, 2 s). **D.** Quantitative analysis of SIC frequency (SCA1, *n* = 8 cells; SCA1+CNO, *n* = 7 cells; Mann-Whitney test; p=0.336). **E.** Representative traces of mI_AHP_ and sI_AHP_ in PCs from SCA1 (red) and SCA1+CNO (dark green) mice (scale bar: 40 pA, 500 ms). **F. Left.** Quantitative analysis of medium I_AHP_ (SCA1, *n* = 10 cells; SCA1+CNO, *n* = 7 cells; t-test, two-tailed p=0.203). mI_AHP_ is measured as the peak current. **Right.** Quantitative analysis of slow I_AHP_ (SCA1, *n* = 10 cells; SCA1+CNO, *n* = 7 cells; t-test, two-tailed p=0.327). sI_AHP_ is measured as the area under the curve during 1 s beginning at the peak of the current. **G.** Cell-attached recordings of spontaneous action potential firing in PCs from SCA1 (red), SCA1+CNO (dark green) mice (scale bar: 100 ms). **H.** Quantitative analysis of PC spontaneous firing rate in the mentioned different conditions (SCA1, *n* = 11 cells; SCA1+CNO, *n* = 13 cells; t-test, two-tailed p=0.151).

## Reference list

Alviña K, Khodakhah K (2010a) The therapeutic mode of action of 4-aminopyridine in cerebellar ataxia. J Neurosci Off J Soc Neurosci 30:7258–7268.

Alviña K, Khodakhah K (2010b) KCa channels as therapeutic targets in episodic ataxia type-2. J Neurosci Off J Soc Neurosci 30:7249–7257.

Anderson CM, Swanson RA (2000) Astrocyte glutamate transport: review of properties, regulation, and physiological functions. Glia 32:1–14.

Andreasen M, Lambert JDC (1995) The excitability of CA1 pyramidal cell dendrites is modulated by a local Ca2+-dependent K+-conductance. Brain Res 698:193–203.

Angulo MC (2004) Glutamate Released from Glial Cells Synchronizes Neuronal Activity in the Hippocampus. J Neurosci 24:6920–6927.

Araque A, Carmignoto G, Haydon PG, Oliet SHR, Robitaille R, Volterra A (2014) Gliotransmitters Travel in Time and Space. Neuron 81:728–739.

Araque A, Parpura V, Sanzgiri RP, Haydon PG (1998) Glutamate-dependent astrocyte modulation of synaptic transmission between cultured hippocampal neurons: Astrocytes modulate synaptic transmission. Eur J Neurosci 10:2129–2142.

Araujo APB, Carpi-Santos R, Gomes FCA (2019) The Role of Astrocytes in the Development of the Cerebellum. The Cerebellum 18:1017–1035.

Beppu K, Kubo N, Matsui K (2021) Glial amplification of synaptic signals. J Physiol 599:2085– 2102.

Bockenhauer D et al. (2009) Epilepsy, Ataxia, Sensorineural Deafness, Tubulopathy, and *KCNJ10* Mutations. N Engl J Med 360:1960–1970.

Borgenheimer E, Hamel K, Sheeler C, Moncada FL, Sbrocco K, Zhang Y, Cvetanovic M (2022) Single nuclei RNA sequencing investigation of the Purkinje cell and glial changes in the cerebellum of transgenic Spinocerebellar ataxia type 1 mice. Front Cell Neurosci 16:998408.

Brockhaus J, Deitmer JW (2002) Long-lasting modulation of synaptic input to Purkinje neurons by Bergmann glia stimulation in rat brain slices. J Physiol 545:581–593.

Buffo A, Rossi F (2013) Origin, lineage and function of cerebellar glia. Prog Neurobiol 109:42– 63.

Burright EN, Brent Clark H, Servadio A, Matilla T, Feddersen RM, Yunis WS, Duvick LA, Zoghbi HY, Orr HT (1995) SCA1 transgenic mice: A model for neurodegeneration caused by an expanded CAG trinucleotide repeat. Cell 82:937–948.

Bushart DD, Chopra R, Singh V, Murphy GG, Wulff H, Shakkottai VG (2018) Targeting potassium channels to treat cerebellar ataxia. Ann Clin Transl Neurol 5:297–314.

Castejón OJ, Dailey ME, Apkarian RP, Castejón HV (2002) Correlative microscopy of cerebellar Bergmann glial cells. J Submicrosc Cytol Pathol 34:131–142.

Cerrato V, Mercurio S, Leto K, Fucà E, Hoxha E, Bottes S, Pagin M, Milanese M, Ngan C-Y, Concina G, Ottolenghi S, Wei C-L, Bonanno G, Pavesi G, Tempia F, Buffo A, Nicolis SK (2018) Sox2 conditional mutation in mouse causes ataxic symptoms, cerebellar vermis hypoplasia, and postnatal defects of Bergmann glia. Glia 66:1929–1946.

Cingolani LA, Gymnopoulos M, Boccaccio A, Stocker M, Pedarzani P (2002) Developmental Regulation of Small-Conductance Ca ^2+^ -Activated K ^+^ Channel Expression and Function in Rat Purkinje Neurons. J Neurosci 22:4456–4467.

Clarke LE, Barres BA (2013) Emerging roles of astrocytes in neural circuit development. Nat Rev Neurosci 14:311–321.

Cohen AS, Coussens CM, Raymond CR, Abraham WC (1999) Long-Lasting Increase in Cellular Excitability Associated With the Priming of LTP Induction in Rat Hippocampus. J Neurophysiol 82:3139–3148.

Custer SK, Garden GA, Gill N, Rueb U, Libby RT, Schultz C, Guyenet SJ, Deller T, Westrum LE, Sopher BL, La Spada AR (2006) Bergmann glia expression of polyglutamine-expanded ataxin-7 produces neurodegeneration by impairing glutamate transport. Nat Neurosci 9:1302– 1311.

Cvetanovic M (2015) Decreased Expression of Glutamate Transporter GLAST in Bergmann Glia Is Associated with the Loss of Purkinje Neurons in the Spinocerebellar Ataxia Type 1. The Cerebellum 14:8–11.

Cvetanovic M, Patel JM, Marti HH, Kini AR, Opal P (2011) Vascular endothelial growth factor ameliorates the ataxic phenotype in a mouse model of spinocerebellar ataxia type 1. Nat Med 17:1445–7.

De Zeeuw CI, Hoebeek FE, Bosman LWJ, Schonewille M, Witter L, Koekkoek SK (2011) Spatiotemporal firing patterns in the cerebellum. Nat Rev Neurosci 12:327–344.

De Zeeuw CI, Hoogland TM (2015) Reappraisal of Bergmann glial cells as modulators of cerebellar circuit function. Front Cell Neurosci 9.

Dell’Orco JM, Wasserman AH, Chopra R, Ingram MAC, Hu Y-S, Singh V, Wulff H, Opal P, Orr HT, Shakkottai VG (2015) Neuronal Atrophy Early in Degenerative Ataxia Is a Compensatory Mechanism to Regulate Membrane Excitability. J Neurosci 35:11292–11307.

Díaz EF, Labra VC, Alvear TF, Mellado LA, Inostroza CA, Oyarzún JE, Salgado N, Quintanilla RA, Orellana JA (2019) Connexin 43 hemichannels and pannexin-1 channels contribute to the α-synuclein-induced dysfunction and death of astrocytes. Glia:glia.23631.

Egorova PA, Gavrilova AV, Bezprozvanny IB (2021) In vivo analysis of the spontaneous firing of cerebellar Purkinje cells in awake transgenic mice that model spinocerebellar ataxia type 2. Cell Calcium 93:102319.

Egorova PA, Zakharova OA, Vlasova OL, Bezprozvanny IB (2016) In vivo analysis of cerebellar Purkinje cell activity in SCA2 transgenic mouse model. J Neurophysiol 115:2840–2851.

Fakhoury M (2018) Microglia and Astrocytes in Alzheimer’s Disease: Implications for Therapy. Curr Neuropharmacol 16:508–518.

Farhy-Tselnicker I, Allen NJ (2018) Astrocytes, neurons, synapses: a tripartite view on cortical circuit development. Neural Develop 13:7.

Fellin T, Pascual O, Gobbo S, Pozzan T, Haydon PG, Carmignoto G (2004) Neuronal Synchrony Mediated by Astrocytic Glutamate through Activation of Extrasynaptic NMDA Receptors. Neuron 43:729–743.

Frisullo G, Della Marca G, Mirabella M, Caggiula M, Broccolini A, Rubino M, Mennuni G, Tonali PA, Batocchi AP (2007) A human anti-neuronal autoantibody against GABAB receptor induces experimental autoimmune agrypnia. Exp Neurol 204:808–818.

Furrer SA, Mohanachandran MS, Waldherr SM, Chang C, Damian VA, Sopher BL, Garden GA, La Spada AR (2011) Spinocerebellar Ataxia Type 7 Cerebellar Disease Requires the Coordinated Action of Mutant Ataxin-7 in Neurons and Glia, and Displays Non-Cell-Autonomous Bergmann Glia Degeneration. J Neurosci 31:16269–16278.

Grewer C, Rauen T (2005) Electrogenic Glutamate Transporters in the CNS: Molecular Mechanism, Pre-steady-state Kinetics, and their Impact on Synaptic Signaling. J Membr Biol 203:1–20.

Grosche J, Matyash V, Möller T, Verkhratsky A, Reichenbach A, Kettenmann H (1999) Microdomains for neuron–glia interaction: parallel fiber signaling to Bergmann glial cells. Nat Neurosci 2:139–143.

Ha GE, Cheong E (2017) Spike Frequency Adaptation in Neurons of the Central Nervous System. Exp Neurobiol 26:179–185.

Halliday GM, Stevens CH (2011) Glia: Initiators and progressors of pathology in Parkinson’s disease: Glia in Parkinson’s Disease. Mov Disord 26:6–17.

Handler HP, Duvick L, Mitchell JS, Cvetanovic M, Reighard M, Soles A, Mather KB, Rainwater O, Serres S, Nichols-Meade T, Coffin SL, You Y, Ruis BL, O’Callaghan B, Henzler C, Zoghbi HY, Orr HT (2023) Decreasing mutant ATXN1 nuclear localization improves a spectrum of SCA1-like phenotypes and brain region transcriptomic profiles. Neuron 111:493–507

Hansen ST, Meera P, Otis TS, Pulst SM (2013) Changes in Purkinje cell firing and gene expression precede behavioral pathology in a mouse model of SCA2. Hum Mol Genet 22:271– 283.

Herson PS, Virk M, Rustay NR, Bond CT, Crabbe JC, Adelman JP, Maylie J (2003) A mouse model of episodic ataxia type-1. Nat Neurosci 6:378–383.

Holt GR, Softky WR, Koch C, Douglas RJ (1996) Comparison of discharge variability in vitro and in vivo in cat visual cortex neurons. J Neurophysiol 75:1806–1814.

Hourez R, Servais L, Orduz D, Gall D, Millard I, de Kerchove d’Exaerde A, Cheron G, Orr HT, Pandolfo M, Schiffmann SN (2011) Aminopyridines Correct Early Dysfunction and Delay Neurodegeneration in a Mouse Model of Spinocerebellar Ataxia Type 1. J Neurosci 31:11795– 11807.

Hoxha E, Balbo I, Miniaci MC, Tempia F (2018) Purkinje Cell Signaling Deficits in Animal Models of Ataxia. Front Synaptic Neurosci 10:6.

Hurlock EC, McMahon A, Joho RH (2008) Purkinje-Cell-Restricted Restoration of Kv3.3 Function Restores Complex Spikes and Rescues Motor Coordination in Kcnc3 Mutants. J Neurosci 28:4640–4648.

Iino M, Goto K, Kakegawa W, Okado H, Sudo M, Ishiuchi S, Miwa A, Takayasu Y, Saito I, Tsuzuki K, Ozawa S (2001) Glia-Synapse Interaction Through Ca ^2+^ -Permeable AMPA Receptors in Bergmann Glia. Science 292:926–929.

Ingram M, Wozniak EAL, Duvick L, Yang R, Bergmann P, Carson R, O’Callaghan B, Zoghbi HY, Henzler C, Orr HT (2016) Cerebellar Transcriptome Profiles of ATXN1 Transgenic Mice Reveal SCA1 Disease Progression and Protection Pathways. Neuron 89:1194–1207.

Irwin S, Vandelft M, Pinchev D, Howell JL, Graczyk J, Orr HT, Truant R (2005) RNA association and nucleocytoplasmic shuttling by ataxin-1. J Cell Sci 118:233–242.

Jayabal S, Chang HHV, Cullen KE, Watt AJ (2016) 4-aminopyridine reverses ataxia and cerebellar firing deficiency in a mouse model of spinocerebellar ataxia type 6. Sci Rep 6:29489.

Jiang R, Diaz-Castro B, Looger LL, Khakh BS (2016) Dysfunctional Calcium and Glutamate Signaling in Striatal Astrocytes from Huntington’s Disease Model Mice. J Neurosci 36:3453– 3470.

Kanai Y, Nussberger S, Romero MF, Boron WF, Hebert SC, Hediger MA (1995) Electrogenic properties of the epithelial and neuronal high affinity glutamate transporter. J Biol Chem 270:16561–16568.

Kasumu AW, Hougaard C, Rode F, Jacobsen TA, Sabatier JM, Eriksen BL, Strøbæk D, Liang X, Egorova P, Vorontsova D, Christophersen P, Rønn LCB, Bezprozvanny I (2012a) Selective positive modulator of calcium-activated potassium channels exerts beneficial effects in a mouse model of spinocerebellar ataxia type 2. Chem Biol 19:1340–1353.

Kasumu AW, Liang X, Egorova P, Vorontsova D, Bezprozvanny I (2012b) Chronic suppression of inositol 1,4,5-triphosphate receptor-mediated calcium signaling in cerebellar purkinje cells alleviates pathological phenotype in spinocerebellar ataxia 2 mice. J Neurosci Off J Soc Neurosci 32:12786–12796.

Khakh BS, Sofroniew MV (2015) Diversity of astrocyte functions and phenotypes in neural circuits. Nat Neurosci 18:942–952.

Kim JH, Lukowicz A, Qu W, Johnson A, Cvetanovic M (2018) Astroglia contribute to the pathogenesis of spinocerebellar ataxia Type 1 (SCA1) in a biphasic, stage-of-disease specific manner. Glia 66:1972–1987.

Kofuji P, Araque A (2021) G-Protein-Coupled Receptors in Astrocyte–Neuron Communication. Neuroscience 456:71–84.

Kofuji P, Newman EA (2004) Potassium buffering in the central nervous system. Neuroscience 129:1043–1054.

Konnerth A, Llano I, Armstrong CM (1990) Synaptic currents in cerebellar Purkinje cells. Proc Natl Acad Sci 87:2662–2665.

Kretzschmar D, Tschäpe J, Bettencourt Da Cruz A, Asan E, Poeck B, Strauss R, Pflugfelder GO (2005) Glial and neuronal expression of polyglutamine proteins induce behavioral changes and aggregate formation in *Drosophila*: Glial Defects by PolyQ Proteins. Glia 49:59–72.

Lin C, Sherathiya VN, Oh MM, Disterhoft JF (2020) Persistent firing in LEC III neurons is differentially modulated by learning and aging. eLife 9:e56816.

Liu D, Nanclares C, Simbriger K, Fang K, Lorsung E, Le N, Amorim IS, Chalkiadaki K, Pathak SS, Li J, Gewirtz JC, Jin VX, Kofuji P, Araque A, Orr HT, Gkogkas CG, Cao R (2022) Autistic-like behavior and cerebellar dysfunction in Bmal1 mutant mice ameliorated by mTORC1 inhibition. Mol Psychiatry.

Magaki SD, Williams CK, Vinters HV (2018) Glial function (and dysfunction) in the normal & ischemic brain. Neuropharmacology 134:218–225.

Maglio LE, Noriega-Prieto JA, Maroto IB, Martin-Cortecero J, Muñoz-Callejas A, Callejo-Móstoles M, Fernández de Sevilla D (2021) IGF-1 facilitates extinction of conditioned fear. eLife 10:e67267.

Mark MD, Krause M, Boele H-J, Kruse W, Pollok S, Kuner T, Dalkara D, Koekkoek S, De Zeeuw CI, Herlitze S (2015) Spinocerebellar ataxia type 6 protein aggregates cause deficits in motor learning and cerebellar plasticity. J Neurosci Off J Soc Neurosci 35:8882–8895.

Massella A, Gusciglio M, D’Intino G, Sivilia S, Ferraro L, Calzà L, Giardino L (2009) Gabapentin treatment improves motor coordination in a mice model of progressive ataxia. Brain Res 1301:135–142.

Matthews EA, Linardakis JM, Disterhoft JF (2009) The Fast and Slow Afterhyperpolarizations Are Differentially Modulated in Hippocampal Neurons by Aging and Learning. J Neurosci 29:4750–4755.

Meunier C, Merienne N, Jollé C, Déglon N, Pellerin L (2016) Astrocytes are key but indirect contributors to the development of the symptomatology and pathophysiology of Huntington’s disease: Astrocytes in Huntington’s Disease. Glia 64:1841–1856.

Milner R ed. (2012) Astrocytes: Methods and Protocols. Chapter 18. Totowa, NJ: Humana Press.

Miyazaki T, Yamasaki M, Hashimoto K, Kohda K, Yuzaki M, Shimamoto K, Tanaka K, Kano M, Watanabe M (2017) Glutamate transporter GLAST controls synaptic wrapping by Bergmann glia and ensures proper wiring of Purkinje cells. Proc Natl Acad Sci 114:7438–7443.

Müller T, Möller T, Neuhaus J, Kettenmann H (1996) Electrical coupling among Bergmann glial cells and its modulation by glutamate receptor activation. Glia 17:274–284.

Nakamura K, Yoshida K, Miyazaki D, Morita H, Ikeda S (2009) Spinocerebellar ataxia type 6 (SCA6): Clinical pilot trial with gabapentin. J Neurol Sci 278:107–111.

Nanclares C, Baraibar AM, Araque A, Kofuji P (2021) Dysregulation of Astrocyte–Neuronal Communication in Alzheimer’s Disease. Int J Mol Sci 22:7887.

Napper RMA, Harvey RJ (1988) Number of parallel fiber synapses on an individual Purkinje cell in the cerebellum of the rat. J Comp Neurol 274:168–177.

Navarrete M, Araque A (2008) Endocannabinoids Mediate Neuron-Astrocyte Communication. Neuron 57:883–893.

Navarrete M, Araque A (2010) Endocannabinoids Potentiate Synaptic Transmission through Stimulation of Astrocytes. Neuron 68:113–126.

Newman E (1993) Inward-rectifying potassium channels in retinal glial (Muller) cells. J Neurosci 13:3333–3345.

Newman EA (1986) High Potassium Conductance in Astrocyte Endfeet. Science 233:453–454.

Newman EA, Frambach DA, Odette LL (1984) Control of Extracellular Potassium Levels by Retinal Glial Cell K ^+^ Siphoning. Science 225:1174–1175.

Nino HE, Noreen HJ, Dubey DP, Resch JA, Namboodiri K, Elston RC, Yunis EJ (1980) A family with hereditary ataxia: HLA typing. Neurology 30:12–20.

Orkand RK, Nicholls JG, Kuffler SW (1966) Effect of nerve impulses on the membrane potential of glial cells in the central nervous system of amphibia. J Neurophysiol 29:788–806.

Paukert M, Agarwal A, Cha J, Doze VA, Kang JU, Bergles DE (2014) Norepinephrine Controls Astroglial Responsiveness to Local Circuit Activity. Neuron 82:1263–1270.

Paulson HL, Shakkottai VG, Clark HB, Orr HT (2017) Polyglutamine spinocerebellar ataxias — from genes to potential treatments. Nat Rev Neurosci 18:613–626.

Perea G, Araque A (2005) Properties of Synaptically Evoked Astrocyte Calcium Signal Reveal Synaptic Information Processing by Astrocytes. J Neurosci 25:2192–2203.

Perea G, Sur M, Araque A (2014) Neuron-glia networks: integral gear of brain function. Front Cell Neurosci 8.

Phillips EC, Croft CL, Kurbatskaya K, O’Neill MJ, Hutton ML, Hanger DP, Garwood CJ, Noble W (2014) Astrocytes and neuroinflammation in Alzheimer’s disease. Biochem Soc Trans 42:1321– 1325.

Piet R, Jahr CE (2007) Glutamatergic and Purinergic Receptor-Mediated Calcium Transients in Bergmann Glial Cells. J Neurosci 27:4027–4035.

Rothstein JD, Dykes-Hoberg M, Pardo CA, Bristol LA, Jin L, Kuncl RW, Kanai Y, Hediger MA, Wang Y, Schielke JP, Welty DF (1996) Knockout of Glutamate Transporters Reveals a Major Role for Astroglial Transport in Excitotoxicity and Clearance of Glutamate. Neuron 16:675–686.

Rudolph R, Jahn HM, Courjaret R, Messemer N, Kirchhoff F, Deitmer JW (2016) The inhibitory input to mouse cerebellar Purkinje cells is reciprocally modulated by Bergmann glial P2Y1 and AMPA receptor signaling: Modulation of Inhibitory Input to Purkinje Cells. Glia 64:1265–1280.

Saab AS, Neumeyer A, Jahn HM, Cupido A, Šimek AAM, Boele H-J, Scheller A, Le Meur K, Götz M, Monyer H, Sprengel R, Rubio ME, Deitmer JW, De Zeeuw CI, Kirchhoff F (2012) Bergmann Glial AMPA Receptors Are Required for Fine Motor Coordination. Science 337:749– 753.

Sah P (1996) Ca2+-activated K+ currents in neurones: types, physiological roles and modulation. Trends Neurosci 19:150–154.

Salinas-Birt A, Zhu X, Lim EY, Cruz Santory AJ, Ye L, Paukert M (2023) Constraints of vigilance-dependent noradrenergic signaling to mouse cerebellar Bergmann glia. Glia:glia.24350.

Santini E, Quirk GJ, Porter JT (2008) Fear Conditioning and Extinction Differentially Modify the Intrinsic Excitability of Infralimbic Neurons. J Neurosci 28:4028–4036.

Sasaki T, Beppu K, Tanaka KF, Fukazawa Y, Shigemoto R, Matsui K (2012) Application of an optogenetic byway for perturbing neuronal activity via glial photostimulation. Proc Natl Acad Sci 109:20720–20725.

Scholl UI, Choi M, Liu T, Ramaekers VT, Häusler MG, Grimmer J, Tobe SW, Farhi A, Nelson-Williams C, Lifton RP (2009) Seizures, sensorineural deafness, ataxia, mental retardation, and electrolyte imbalance (SeSAME syndrome) caused by mutations in *KCNJ10*. Proc Natl Acad Sci 106:5842–5847.

Schousboe A (2000) Pharmacological and functional characterization of astrocytic GABA transport: a short review. Neurochem Res 25:1241–1244.

Seidel K, Siswanto S, Brunt ERP, den Dunnen W, Korf H-W, Rüb U (2012) Brain pathology of spinocerebellar ataxias. Acta Neuropathol (Berl) 124:1–21.

Seki T, Sato M, Kibe Y, Ohta T, Oshima M, Konno A, Hirai H, Kurauchi Y, Hisatsune A, Katsuki H (2018) Lysosomal dysfunction and early glial activation are involved in the pathogenesis of spinocerebellar ataxia type 21 caused by mutant transmembrane protein 240. Neurobiol Dis 120:34–50.

Serra HG, Byam CE, Lande JD, Tousey SK, Zoghbi HY, Orr HT (2004) Gene profiling links SCA1 pathophysiology to glutamate signaling in Purkinje cells of transgenic mice. Hum Mol Genet 13:2535–43

Servadio A, Koshy B, Armstrong D, Antalffy B, Orr HT, Zoghbi HY (1995) Expression analysis of the ataxin–1 protein in tissues from normal and spinocerebellar ataxia type 1 individuals. Nat Genet 10:94–98.

Shakkottai VG, do Carmo Costa M, Dell’Orco JM, Sankaranarayanan A, Wulff H, Paulson HL (2011) Early changes in cerebellar physiology accompany motor dysfunction in the polyglutamine disease spinocerebellar ataxia type 3. J Neurosci Off J Soc Neurosci 31:13002–13014.

Sheeler C, Rosa J-G, Borgenheimer E, Mellesmoen A, Rainwater O, Cvetanovic M (2021) Post-symptomatic Delivery of Brain-Derived Neurotrophic Factor (BDNF) Ameliorates Spinocerebellar Ataxia Type 1 (SCA1) Pathogenesis. The Cerebellum 20:420–429.

Shim HG, Jang S-S, Kim SH, Hwang EM, Min JO, Kim HY, Kim YS, Ryu C, Chung G, Kim Y, Yoon B-E, Kim SJ (2018) TNF-α increases the intrinsic excitability of cerebellar Purkinje cells through elevating glutamate release in Bergmann Glia. Sci Rep 8:11589.

Shimamoto K, Lebrun B, Yasuda-Kamatani Y, Sakaitani M, Shigeri Y, Yumoto N, Nakajima T (1998) dl- *threo* -β-Benzyloxyaspartate, A Potent Blocker of Excitatory Amino Acid Transporters. Mol Pharmacol 53:195–201.

Shin J-Y, Fang Z-H, Yu Z-X, Wang C-E, Li S-H, Li X-J (2005) Expression of mutant huntingtin in glial cells contributes to neuronal excitotoxicity. J Cell Biol 171:1001–1012.

Shuvaev AN, Belozor OS, Mozhei O, Yakovleva DA, Potapenko IV, Shuvaev AN, Smolnikova MV, Salmin VV, Salmina AB, Hirai H, Teschemacher AG, Kasparov S (2021) Chronic optogenetic stimulation of Bergman glia leads to dysfunction of EAAT1 and Purkinje cell death, mimicking the events caused by expression of pathogenic ataxin-1. Neurobiol Dis 154:105340.

Sloan SA, Barres BA (2014) Mechanisms of astrocyte development and their contributions to neurodevelopmental disorders. Curr Opin Neurobiol 27:75–81.

Sofroniew MV, Vinters HV (2010) Astrocytes: biology and pathology. Acta Neuropathol (Berl) 119:7–35.

Storm JF (1987) Action potential repolarization and a fast after-hyperpolarization in rat hippocampal pyramidal cells. J Physiol 385:733–759.

Sugihara I (2005) Microzonal projection and climbing fiber remodeling in single olivocerebellar axons of newborn rats at postnatal days 4-7. J Comp Neurol 487:93–106.

Tsai PT, Hull C, Chu Y, Greene-Colozzi E, Sadowski AR, Leech JM, Steinberg J, Crawley JN, Regehr WG, Sahin M (2012) Autistic-like behaviour and cerebellar dysfunction in Purkinje cell Tsc1 mutant mice. Nature 488:647–651.

Vianello M, Tavolato B, Armani M, Giometto B (2003) Cerebellar ataxia associated with anti-glutamic acid decarboxylase autoantibodies. The Cerebellum 2:77–79.

Volterra A, Meldolesi J (2005) Astrocytes, from brain glue to communication elements: the revolution continues. Nat Rev Neurosci 6:626–640.

Voogd J, Glickstein M (1998) The anatomy of the cerebellum. Trends Neurosci 21:370–375.

Wang F, Xu Q, Wang W, Takano T, Nedergaard M (2012) Bergmann glia modulate cerebellar Purkinje cell bistability via Ca ^2+^ -dependent K ^+^ uptake. Proc Natl Acad Sci 109:7911–7916.

Wu H-S, Sugihara I, Shinoda Y (1999) Projection patterns of single mossy fibers originating from the lateral reticular nucleus in the rat cerebellar cortex and nuclei. J Comp Neurol 411:97– 118.

Wulff P, Goetz T, Leppä E, Linden A-M, Renzi M, Swinny JD, Vekovischeva OY, Sieghart W, Somogyi P, Korpi ER, Farrant M, Wisden W (2007) From synapse to behavior: rapid modulation of defined neuronal types with engineered GABAA receptors. Nat Neurosci 10:923–929.

Wulff P, Schonewille M, Renzi M, Viltono L, Sassoè-Pognetto M, Badura A, Gao Z, Hoebeek FE, van Dorp S, Wisden W, Farrant M, De Zeeuw CI (2009) Synaptic inhibition of Purkinje cells mediates consolidation of vestibulo-cerebellar motor learning. Nat Neurosci 12:1042–1049.

Yamada K, Watanabe M (2002) Cytodifferentiation of Bergmann glia and its relationship with Purkinje cells. Anat Sci Int 77:94–108.

Yamanaka K, Chun SJ, Boillee S, Fujimori-Tonou N, Yamashita H, Gutmann DH, Takahashi R, Misawa H, Cleveland DW (2008) Astrocytes as determinants of disease progression in inherited amyotrophic lateral sclerosis. Nat Neurosci 11:251–253.

Zerangue N, Kavanaugh MP (1996) Flux coupling in a neuronal glutamate transporter. Nature 383:634–637.

